# Combining quadrat, rake and echosounding to estimate submerged aquatic vegetation biomass at the ecosystem scale

**DOI:** 10.1101/2022.03.15.484486

**Authors:** Morgan Botrel, Christiane Hudon, Pascale M. Biron, Roxane Maranger

## Abstract

Measuring freshwater submerged aquatic (SAV) biomass at large spatial scales is challenging and no single technique can cost effectively accomplish this while maintaining accuracy. We propose to combine and intercalibrate accurate quadrat-scuba diver technique, fast rake sampling and large scale echosounding. We found that the relationship between quadrat and rake biomass is moderately strong (R^2^ = 0.62, RMSECV = 2.19 g/m_2_) and varies with substrate type and SAV growth form. Rake biomass was also successfully estimated from biovolume_10_ and its error (R^2^ = 0.53, RMSECV = 5.95 g/m^2^), a biomass proxy derived from echosounding, at a resolution of 10 m radius from rake sampling point. However, the relationship was affected by SAV growth form, depth, acoustic data quality and wind conditions. Sequential application of calibrations yielded predictions in agreement with quadrat observations, but echosounding predictions underestimated biomass in shallow areas (< 1.5 m) while outperforming point estimation in deep areas (> 3 m). Whole-system biomass was more accurately estimated by calibrated echosounding than rake point surveys, owing to the large sample size and better representation of spatial heterogeneity of echosounding. We recommend developing as a one-time event a series of quadrat and rake calibration equations for each growth form and substrate type. Because the relationship between biovolume and biomass depends on SAV growth form, rake and echosounding calibration needs to be conducted frequently. With the two calibrations, rake can thus be used as a rapid ground truthing or in shallow areas where echosounding is inadequate.

## Introduction

Submerged aquatic vegetation (SAV) provides many aquatic ecosystem functions and services, from stabilizing sediments to maintaining critical habitat for fauna (Caraco et al. 2006; Carpenter and Lodge 1986; Hilt et al. 2017). Ecosystem service provisioning by SAV meadows is dependent both on plant patch density and size where high elemental fluxes and faunal populations are associated with high SAV standing stock (e.g. Brown et al. 2004; Cyr and Downing 1988; Rooney et al. 2003). However, SAV standing stock is sensitive to human pressures. For example, declining SAV abundance in shallow lakes mainly reflects loss of water transparency as a function of increasing eutrophication (Scheffer et al. 2001; Scheffer et al. 1993). Management efforts around SAV often attempt to restore abundant meadows, however invasive alien aquatic ‘weeds’ that impede multiple water usages are typically actively removed (Hussner et al. 2017; Madsen and Wersal 2017). Regardless of the management needs, accurate estimates of SAV standing stock are essential to assess their overall functional role in ecosystems and whether management strategies are working.

SAV standing stock or standing biomass is the density measure of aboveground living plant material in mass per unit area. Biomass can be either measured using destructive removal techniques or estimated using remote sensing (Figure 1a). Destructive techniques consist of harvesting plant material, either from below or above the water surface. These are measures of direct biomass, as opposed to a proxy, and represent SAV biomass spatially as a point phenomenon. Direct underwater harvest of SAV by scuba divers is the most accurate biomass measure and can be used at all depths (Downing and Anderson 1985; Madsen 1993). However, this technique requires specialized scuba training and, because of long collection time, small quadrat size (< 1 m^2^), and the associated safety risks and expenses, all of which hinder its use for extensive sampling or for sampling in turbid waters. To alleviate these shortcomings, destructive techniques using tools to grab samples from the surface have been developed. Multiple tools exist, but the double-headed rake with a variety of assembly features (on a telescopic pole, tongs or rope) and collection methods (dragged, griped or spun) is rather well adapted for use in large surveys (Johnson and Newman 2011; Madsen and Wersal 2017; Rodusky et al. 2005; Yin and Kreiling 2011). Indeed, the rake is convenient for large-scale recurrent SAV sampling due to its low purchase cost, ease of use, fast collection, robustness and simple maintenance. However, rake collection is restricted to shallow depths (< 3 - 4 m) and is biased since plant material is often not entirely collected or collected in excess (Johnson and Newman 2011; Kenow et al. 2007; Rodusky et al. 2005). Furthermore, all destructive methods are point measurements and require the collection and processing of multiple replicates to reach a reasonably precise biomass estimate due to the inherent patchiness of SAV meadows (Downing and Anderson 1985). Accurate measurements of SAV biomass capturing SAV heterogeneity at large spatial scales thus remain a challenge, since surveys using quadrats are only accurate at a very small scale whereas rake samples are biased but provide a broader estimate of biomass distribution.

**Figure 1:**
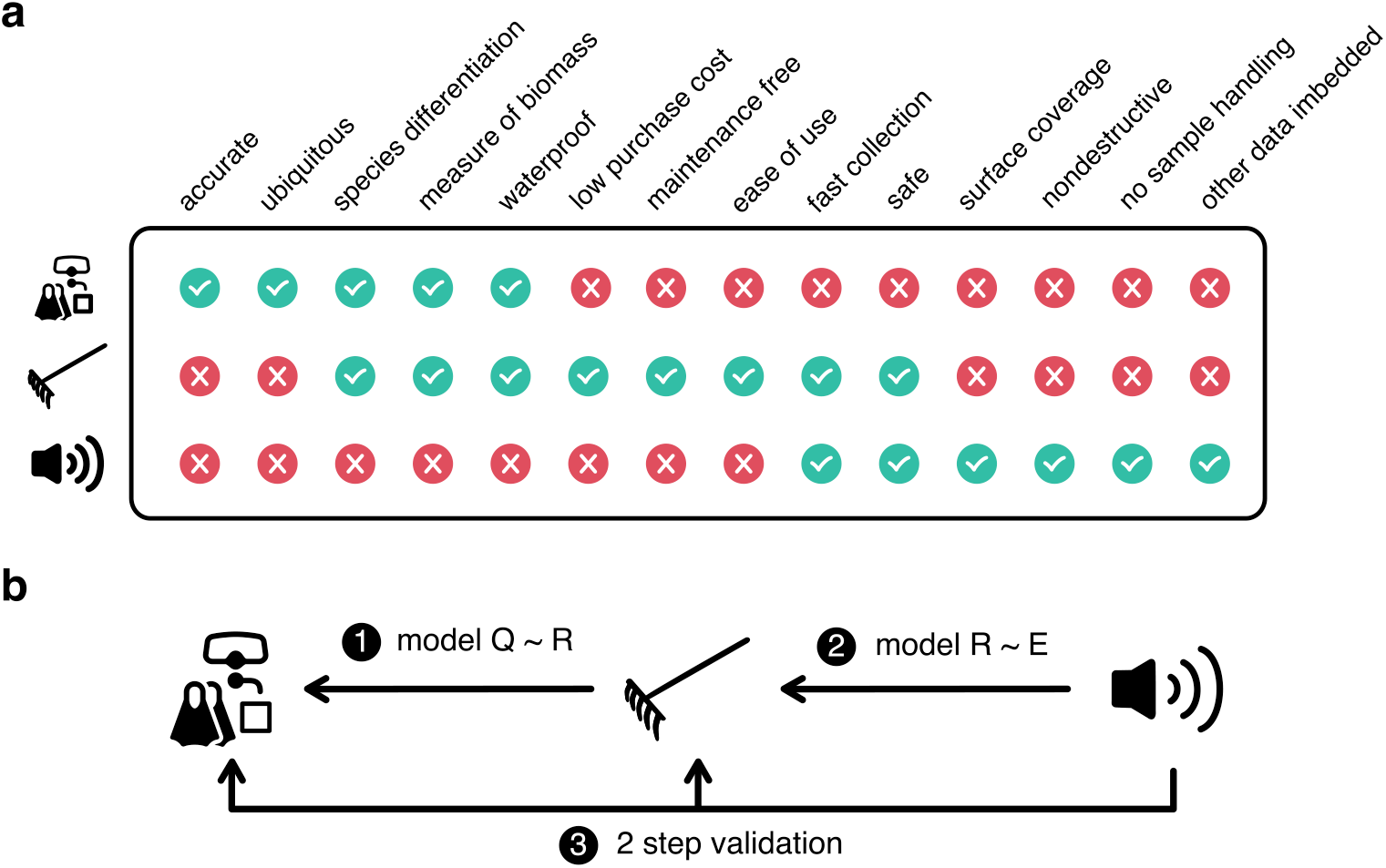
Comparison and combination of the quadrat, rake and echosounding techniques to estimate submerged aquatic vegetation biomass. A) Strengths and weaknesses of the different techniques. B) Approach and steps to combine techniques. Q: quadrat, R: rake.

At large spatial scales, SAV biomass estimates can be improved with remote sensing that represent SAV in space as a continuous surface phenomenon. These techniques provide a proxy of biomass where SAV is detected by a receiver that captures a signal (e.g. sound or light) reflected or emitted by SAV (Rowan and Kalacska 2021). Capturing the sound reflection made by gas vacuoles in plant tissue (the echo) using an active single beam echosounder, is the simplest and best remote sensing technique to estimate SAV biomass in turbid freshwaters (Duarte 1987; Rowan and Kalacska 2021; Sabol et al. 2002; Vis et al. 2003). Indeed, echosounders are transportable and increasingly affordable devices that, just like a sonar used for fishing, can be hooked on the side of a boat for a rapid, non-destructive and repeatable survey (e.g. (Howell and Richardson 2019). The reflection of sound on bottom surfaces and canopies allows for the simultaneous measurement of SAV height and water depth, which is extremely useful as water depth strongly influences SAV biomass (Duarte and Kalff 1990). Because SAV height correlates to biomass, echosounding has successfully been used to model whole community biomass (Duarte 1987; Maceina et al. 1984; Sabol et al. 2002). However, the allometric relationship between height and biomass varies with species growth form and thus the calibration of SAV echo is species- or stand-specific and as such the measurement needs to be repeated frequently (Duarte 1987). Other drawbacks of echosounding include its lack of species differentiation, inability to take measurements in very shallow waters (< 0.4 – 0.7 m) or when plants reach water surface, and obligate sampling during calm weather conditions, since wind-induced gas bubbles strongly scatter sound (Sabol et al. 2002). Furthermore, the use of echosounding requires technical expertise, from maneuvering the instrument, electronic maintenance to processing data output.

Each of the aforementioned techniques has several advantages and disadvantages for estimating SAV biomass depending on the scale of study (Figure 1a). One promising approach would be to design a process enabling the benefits of all three techniques which could provide accurate biomass estimates that take into account SAV heterogeneity at broad spatial scales. Therefore, our objective is to develop a process that assesses the interchangeability of quadrat, rake and echosounding techniques to render their use synergistic and cost-effective. To do so, we conducted two intercalibrations: one between biomass from quadrat and rake and the other between rake and a biomass proxy derived from echosounding (Figure 1b). Furthermore, we investigated how the predictions of both intercalibrations models are affected by environmental factors. The resulting two models were then sequentially applied to echosounding data to validate our approach. Finally, we assessed how our approach modifies whole-system biomass estimation.

## Materials and Procedures

To compare, and integrate the different biomass assessment methods, three datasets were used: the quadrat-rake, the rake-echosounding and the validation datasets (Table 1). All datasets were collected in Lake Saint-Pierre (LSP), a ∼300 km^2^ widening of the Saint-Lawrence River in Quebec, Canada (Figure 2a, b). Part of the quadrat and rake dataset was also collected in the nearby upstream fluvial lakes, Saint-Louis and Saint-François.

**Figure 2:**
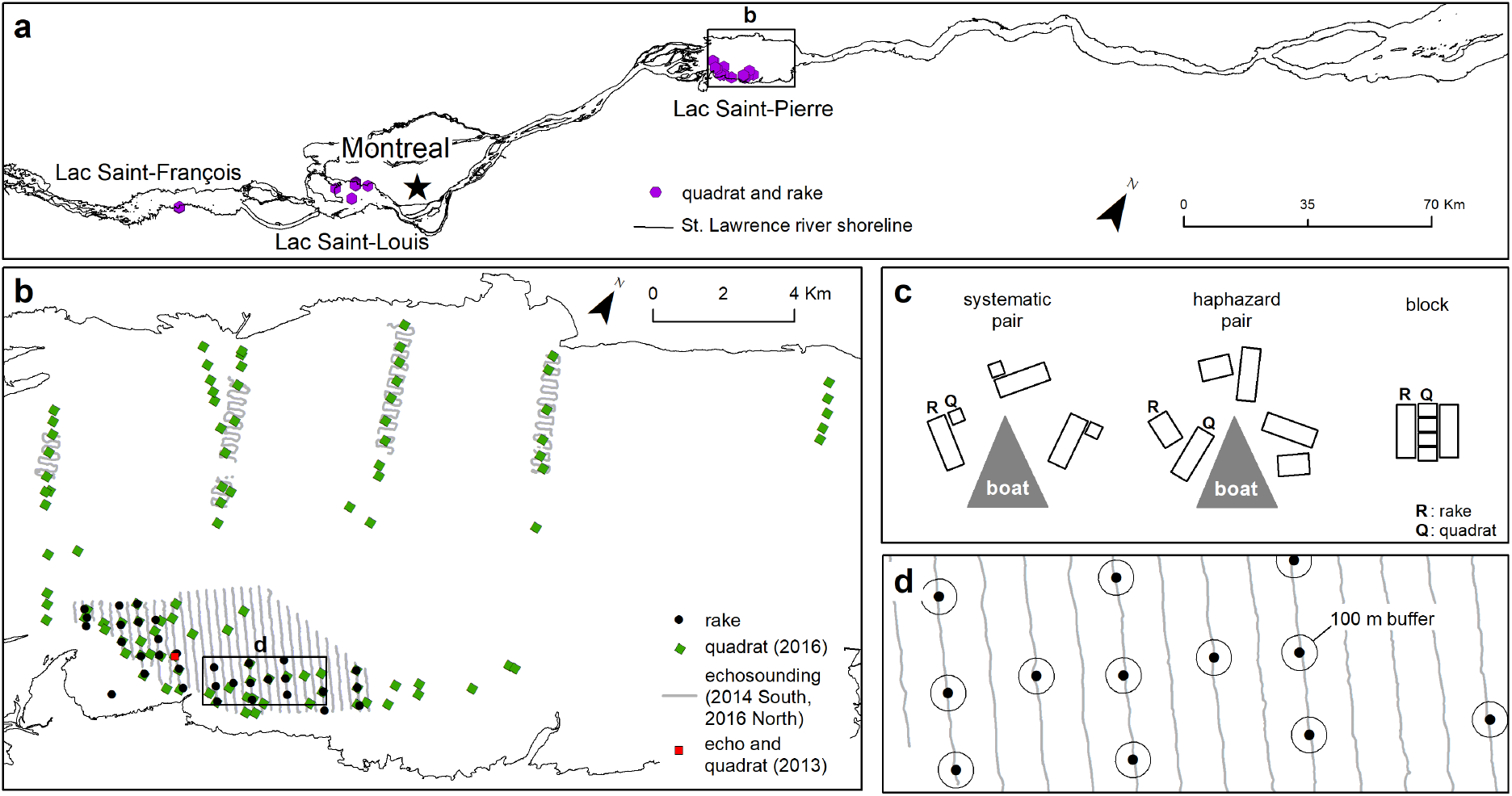
a) Sampling locations in the St. Lawrence River for the quadrat-rake (Q-R) dataset; b) rake-echosounding (R-E) and validation (echosounding - quadrat) datasets in Lac Saint-Pierre. C) collection strategy to compare quadrat to rake; d) collection strategy to compare rake to echosounding. For the R-E dataset, 2014 rake sampling sites and echosounding tracks are shown as an example (South-West sector in b). To clearly visualize the validation dataset, all the 2016 quadrats (green diamonds in b) are depicted but only part of the 2016 echosounding tracks (North section in b, South-West tracks are similar to R-E dataset) and the general location of the 2013 small scale sampling (red square, South-West sector in b) are shown.

**Table 1:**
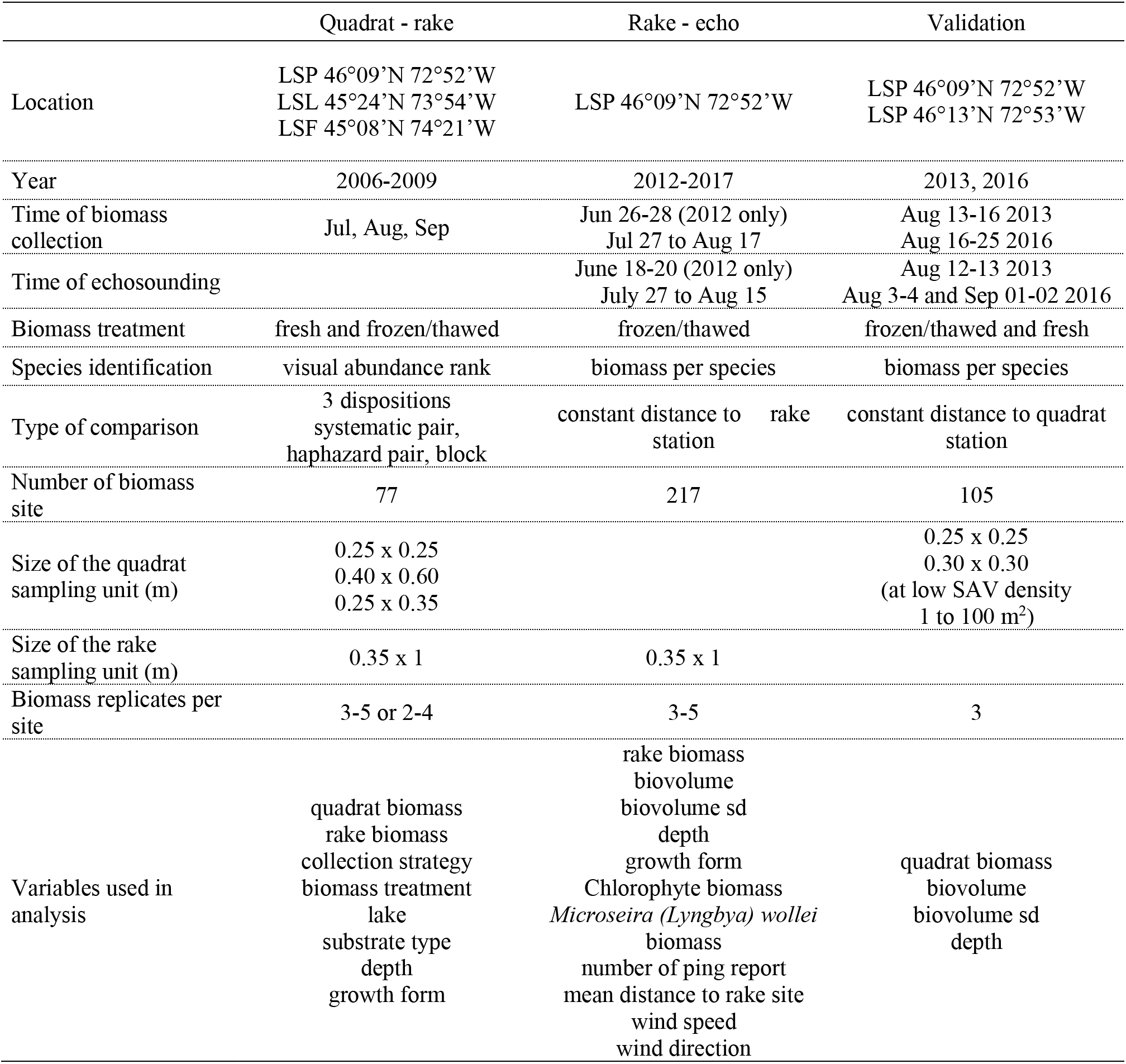
Summary of the three datasets used in this study. LSP Lake Saint-Pierre, LSL Lake Saint-Louis, LSF Lake Saint-François, sd standard deviation.

### Quadrat-rake (Q-R) dataset

The first dataset was used to assess the correspondence between biomass estimates derived from quadrat and rake samples (hereafter named the Q-R dataset), using one of three different collection strategies: systematic pairs, haphazard pairs or blocks (Figure 2c). For the first two strategies, paired rake and quadrat samples were collected around the anchored boat, using either a systematic or a haphazard strategy. In the “systematic pair” strategy, a single quadrat (0.25 m x 0.25 m) was systematically collected to the upper right side of each rake sample (1 m x 0.35 m). In the “haphazard pair” strategy, bigger quadrats (0.40 m x 0.60 m) were located in the vicinity of the rake sample site. In the block sampling strategy, a block, which consisted of two rake samples positioned on both sides of a row of 4 individual quadrats (0.25 m x 0.35 m each) aligned to mimic the raked area, was collected on each side of the boat.

Quadrat plant samples were harvested by divers from within a PVC frame placed on the lake bottom by divers. All aboveground plant material was cut using grass-clippers or broken at the sediment surface. Rake samples were collected from the anchored boat, using a double-headed rake (0.35-m head width with fourteen, 8-cm long teeth on both sides) mounted on a telescopic pole (maximum length 5 m) following the method described in Yin and Kreiling (2011). The rake was lowered in the water and dragged toward the boat over the bottom on a length of approximately 1 m. As it was lifted from the water, the rake was flipped 180° to minimize plant loss.

Biomass samples were collected during the period of maximum SAV development during the summers of 2006 to 2009. Sites were chosen to cover a wide range of water depths, sediment types and SAV biomass. Water depth at each site was measured with a survey ruler and sediment type was classified as pebble, sand, silt, clay or a mixture thereof. SAV species composition of each sample was visually assessed in decreasing abundance rank (1 = most abundant species). Plant material was washed on site to remove sediment and debris prior to further processing (on site, in the laboratory on fresh or thawed samples). Macrophytes (vascular plants and macroalgae such as *Nitella* and *Chara* spp) were separated from filamentous algal mats (Chlorophytes or the cyanobacterium *Microseira* (*Lyngbya) wollei*; each group was wrung out manually and either weighted with a hook-scale on site (precision 0.02 kg) or an electronic scale in the laboratory (precision 0.1 g). A subsample of filamentous algae was preserved in lugol for subsequent microscopic identification (250 X). Wet mass was converted to dry mass using previously established conversion factors for SAV (Hudon et al. 2012) and filamentous species (Cattaneo et al. 2013). All of the manipulations from sample collection to final biomass conversion are referred to ‘biomass treatments’.

Both biomass and species information were aggregated per site. Biomass measurements were reported as a mean of 3-5 rake and quadrat replicates for the systematic and haphazard pair collection strategies or on 2 rakes and 4 quadrats for the block collection strategy. Species information per replicate sample was converted to dominant growth form type as either species forming dense canopy, short understory species or species with leaves growing from a basal rosette, using the dominant species only (rank 1). Canopy-forming plants (< 3 m) included mainly *Potamogeton richardsonii* but also *Heteranthera dubia*, *Stuckenia pectinata*, *Elodea* spp and *Myriophyllum* spp. *Chara* spp formed a low-lying (< 20 cm) layer on the bottom, while *Vallisneria americana* formed rosette of linear leaves extending towards the surface (< 1.5 m). When more than half of the replicates at a given site had the same dominant growth form, that growth form was allocated to the site.

### Rake and echosounding (R-E) dataset

The second dataset was used to predict rake biomass from echosounding (hereafter named R-E dataset). Both acoustic and rake surveys were carried out in the southwest portion of LSP once a year from 2012 to 2017 during the moment of maximum biomass, but also once early in the growing season in June 2012 (Figure 2b, d). Acoustic surveys were conducted on 250 m-spaced transects perpendicular to the lake shore at low and constant speed (∼1.1 m/sec). Acoustic data were collected using a downward-looking BioSonics single beam transducer mounted on an Ocean Science riverboat fixed to the side of the vessel by a steel rod. This set-up allowed the transducer to be constantly immersed below 5 cm of the water surface. Floating and drifting vegetation frequently became entangled with the transducer, which consequently was regularly cleaned. The transducer had a beam angle of 6.6° and a working frequency of 430 kH, which depending on water depth, represents a circle of 0.10 to 0.20 m cone diameter of sound reaching the sediment surface. Echosounding was controlled by a BioSonics DTX system running Visual Acquisition 6.06 with a pulse length of 0.1 msec and a ping rate of 5 ping/sec. Geolocation data were simultaneously recorded with a GNSS NovAtel Smart V1 receiver placed on top of the transducer. Real-time differential correction was obtained using the Omnistar VBS network in 2012 and 2013 (0.90 m precision) and the WAAS network in 2014 to 2017 (0.65 m precision). Acoustic data (echograms and geographic coordinates) were saved in dt4 files on a laptop PC for post-processing.

Acoustic data were processed using Visual Habitat 1. Lake bottom was first determined using a rising edge threshold of -47 dB and a rising edge length criterion of 10 cm. The resulting delineations on the echograms were manually corrected to have a bottom line following highest amplitudes. SAV was subsequently analyzed using plant detection threshold above ambient noise of -68 dB and a minimum height of plant detection of 10 cm. SAV delineations were again manually corrected when the plant line was going beneath the bottom line, mainly due to false detection of plants reaching the water surface. Invalid ping reports were removed, and remaining reports were exported for cycles of 5 pings to have a sample resolution similar to the raked length and a data point every ∼1.1 m. For each cycle, mean plant height (m), plant cover (%), mean geographic position (dd.dddd) and mean depth (m) were exported. SAV biovolume, a potential proxy of SAV biomass, was computed as mean plant height × mean plant cover.

Within one to two week of the acoustic surveys, rake samples were collected at 35 sites along the echosounding transects. Sites were positioned using a Trimble GeoXT (precision 0.50 m) in 2012-2013, a SXBlue II GPS (precision 0.65 m) in 2014-2015 and a Garmin 64S (precision 3 m) in 2016 and 2017. Water depth (z) was measured with a survey ruler and, to be comparable with depth during echosounding, was corrected using the equation z = z_rake_ – lvl_date rake_ + lvl_date echo_ and water level (lvl) at station 15975 (Fisheries and Ocean Canada, www.isdm-gdsi.ca, accessed 2017/11/27). At each site, 3-5 rake replicates were collected around the anchored, 7-m long boat, using the same apparatus and dragged length as the Q-R dataset. To ensure that a location was not raked twice, rakes were systematically collected in front and on each sides of the boat and as distant as possible when replicate number exceeded three. Plant material was similarly processed as described for the Q-E dataset, but plants were sorted by species in the lab and macrophyte biomass was measured on dried material. Total SAV biomass was computed from the species biomass and pooled by growth form. The same species as the Q-R dataset were found. All SAV biomass herein are reported as mean dry biomass in g/m^2^.

### Echosounding and quadrat validation dataset

The third dataset was used to validate biomass estimation from echosounding and compare it to quadrat biomass (hereafter named the validation dataset). Acoustic and quadrat surveys were carried at the time of maximum biomass accumulation in 2013 and in 2016. In 2013, the survey was concentrated in a small (100 m x 100 m) area of high biomass in southwest LSP characterized by a narrow depth range (1.4 - 1.5m, Figure 2b). In 2016, the survey was conducted in both the southwest and the northern sector of LSP. The northern sector was characterized by greater depth range (0.5 - 7.0 m) due to the crossing of the St. Lawrence navigation channel. Quadrat sampling in this sector was carried in 5 transects and the 250 m-spaced echosounding transects were perpendicular to the quadrat transects. The southwest sector had shallower depths (< 2.7 m) and echosounding transects were located following the same methodology as described for the R-E datasets. Processing of acoustic data and biomass samples were also conducted similarly to the R-E dataset, but aboveground biomass was collected in triplicates by scuba divers using a 0.30 m x 0.30 m quadrat in 2013, a 0.25 m x 0.25 m quadrat in 2016 and balance precision was 0.0004 g. When SAV density was very low, divers evaluated the surface they patrolled without plants as a straight line or a circle around the anchor (up to 100 m^2^) and collected the few plants they found.

### Matching rake with echosounding and determining optimal spatial resolution

To assess the correspondence between rake biomass and the biomass proxy from echosounding, we first needed to spatially match rake site to the echosounding track and, second, determine the spatial resolution of the aggregated rake replicates to which we could compare the echosounding data. Our approach was to calculate the mean biovolume of the acoustic data falling in a circle at increasing radial distance from the rake site, ranging from 1 to 100 m (Figure 2d). Higher correlations between the biovolume_D_ calculated at a given distance (D) and rake biomass were interpreted as closer to the resolution of rake collection. To do so, we first imported ping reports and biomass site positions transformed in UTM 18 in QGIS (QGIS Development Team). We computed a 100 m-radius buffer around each biomass site and echosounding track data were intersected with the buffers. Using the intersected layer, all geographic distances between the individual ping reports and the nearest biomass sites (i.e. the target layer) were calculated using the distance matrix tool. In R and using the intersected table, we then calculated mean biovolume_D_ per radial distance, at 1m increments between 1 and 40 m and at 5 m increments between 40 and 100 m. This set-up also allowed us to evaluate the effect of acoustic data quality (error, proximity, and frequency) on biomass prediction. We calculated the biovolume_D_ standard deviation (sd) as an indicator of the acoustic error, the mean ping report distance to rake site as the acoustic proximity, and the number of ping reports per site as the acoustic frequency. A similar method was applied to the validation dataset to derive biomass comparable acoustic data, but only using the rake spatial resolution that was determined from the R-E dataset statistical analysis.

### Statistical analysis

To predict quadrat biomass from rake biomass, ordinary least square regression (OLS) with leave-one-out cross-validation was used. Since variables describing our sampling conditions were mostly categorical (collection strategy, biomass treatment, lake, substrate type, growth form, Table 1), we tested their effect on the quadrat-rake biomass relationship using ANCOVA. When the relations between quadrat and rake biomass exhibited different slopes and intercepts for a given condition, separate regression equations were computed for each category using OLS. We also evaluated the effect of depth on the quadrat-rake biomass relationship using partial regression.

To estimate the optimal spatial resolution to compare rake and echosounding, we used Pearson correlation analysis between rake biomass and the average biovolume at various distances from the biomass site (biovolume_D_). Using the dataset with the determined resolution, we evaluated how the sampling conditions varied between field campaigns by applying analysis of variance or Kruskal-Wallis test when variance was not homogeneous across groups (previously analyzed using Bartlett test). Differences among groups were assessed using post-hoc parametric or non-parametric test, when appropriate (t-test or Wilcoxon test), with Holm correction for multiple tests. Variables included in the analysis described SAV abundance, SAV growth form, macroalgae abundance, water depth, wind (hourly wind speed and direction acquired from Meteorological Service of Canada, climate.weather.gc.ca, accessed 2017/11/27) and acoustic data quality during surveys (Table 1). We then fitted rake biomass to the multiple potential predictors using partial least square regression (PLSR). This method is robust when there is high collinearity among many predictors and when numbers of observations are low (Mevik and Wehrens 2007). We first selected components using the leave-one-out cross-validation and the one-sigma heuristic approach. We then selected variables using the filter method of selectivity ratio (SR), which is the ratio of the explained to the residual variance of the X variables on the y target projection. We chose SR over the commonly used variable importance of the projection (VIP) because the former selects important variables using a F-test and performs well for prediction (Farrés et al. 2015).

To validate our intercalibration approach, we applied our models in two steps. First, from echosounding biovolume_D_, we predicted rake biomass and from that estimation we predicted quadrat biomass. We propagated model error with Monte Carlo simulations and used the root mean square error of cross-validation (RMSECV) as the model error term. Second, we visually compared in space the quadrat-equivalent biomass predicted from echosounding to measured quadrat biomass. For this, we created spatially interpolated biomass maps from both quadrat and echosounding using kriging.

To compare the effect of sample size and spatial coverage on whole-system biomass, we compared biomass estimate from point sampling to remote sensing using the R-E dataset over five summer campaigns. We first calculated mean biovolume_D_ along echosounding transects in distance bins corresponding to the determined resolution. We then created for each campaign a spatial polygon where both rake and echosounding had a good spatial coverage. To do so, we intersected for each campaign two 100 m buffers around concave hulls created from the rake sites and echosounding sites. Within the intersected spatial polygon, mean biomass for rake and echosounding sites were then calculated with bootstrapping confidence intervals. This resampling technique has been shown to be more appropriate for estimating seaweed confidence interval and is particularly suitable when the true distribution of the data is unknown or skewed (Johnson 2020). For echosounding, we also estimated a spatially interpolated average.

Due to high SAV heterogeneity ranging from bare patches to dense plant abundance, absence data (where quadrat or rake biomass = 0 and biovolume = 0) were excluded from modelling analysis. The three techniques had different sample unit size and sampling effort (Table 1). Therefore, techniques with higher sample unit size, e.g. rake, or having a higher number of observations, e.g. echosounding, had a higher probability of finding SAV in low biomass regions compared to their correspondent explained variable (e.g. quadrat and rake respectively). The resulting models to predict rake and quadrat biomass were applied only to presence data (biovolume_D_ > 0 or rake biomass > 0) and absence recorded by both techniques were deemed true. When necessary, data used in statistical analysis were transformed to respect assumptions of normality. Models were evaluated for their ability to predict biomass by minimizing the RMSECV and the mean absolute percentage error (MAPE) of cross-validation predictions. Data handling and statistical analyses were performed in R 3.6.3 (R core team 2020). Cross validation for OLS was computed using the package caret and PLSR with variable selection carried out using pls and plsVarSel (Kuhn 2020, Mevik et al. 2020, Mehmood et al. 2012). Geospatial data was handled using the sp, sf, raster and concaveman packages (Pebesma and Bivand 2005, Pebesma 2018, Hijmans 2020, Gombin et al. 2020), while geostatistical modelling and kriging was performed using gstat (Pebesma 2004). Bootstrapping confidence intervals were computed using the boot package (Canty and Ripley 2021).

## Assessment

### Prediction of quadrat biomass using rake samples

The overall relationship between quadrat biomass and rake biomass was moderately strong with an R^2^ of 0.62 (Figure 3a, Table 2). The quadrat and rake distributions were skewed, leading to non-normal regression residuals, therefore the modelled relationship was on log_10_ data. The quadrat biomass was generally significantly higher by a factor of 4 than biomass estimated using a rake (median_Q_ = 55.19 g/m^2^, median_R_ = 13.83 g/m^2^, paired *t*-test *p* < 0.0001). Quadrat estimates had 20 times higher variance (median s^2^_R_ = 536.60 g/m^2^, median s^2^_Q_ = 29.79 g/m^2^, *p* < 0.001) than rake equivalents, resulting in a higher standard error (SE, median SE_Q_ = 10.48 g/m^2^, median SE_R_ = 3.57 g/m^2^). There was a correlation between mean biomass and its associated error for both quadrat and rake (*r*_Q_ = 0.82, *r*_R_ = 0.77), indicating that the difference in error between methods could be caused by the higher biomass measured in quadrat sampling. Furthermore, this inflation of error with biomass was not constant: the ratio of variance to mean biomass (s^2^/*x̄*) increased with mean biomass and was generally higher for quadrat than rake (*x̄*_Q_ = 41, *x̄*_R_ = 17). As a result, both rake and quadrat measurements showed a spatial aggregation of biomass (s^2^ > *x̄*), although more markedly so for quadrats (99% of observations) than for rakes (67% of observations).

**Figure 3:**
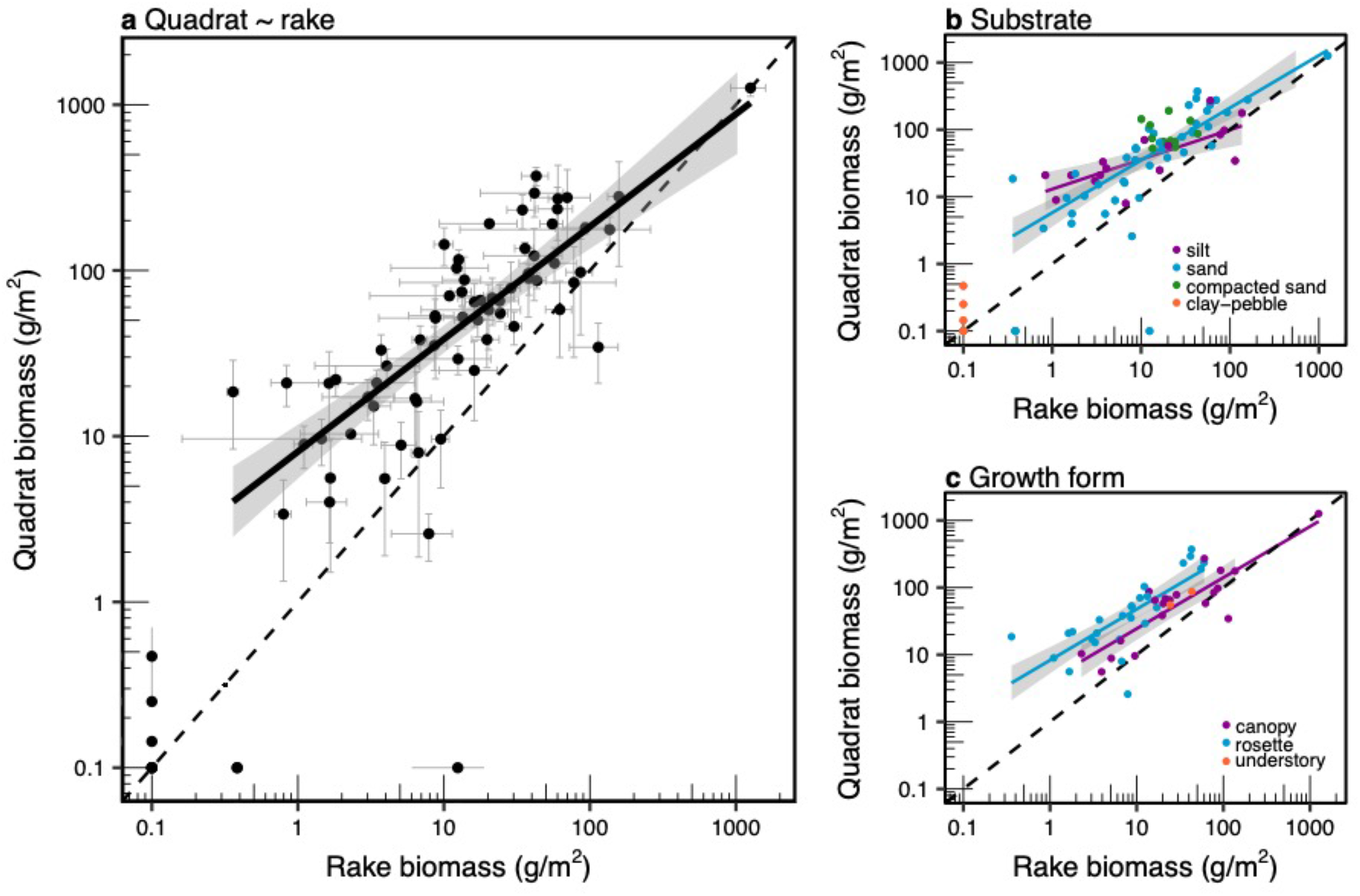
Relationships to predict quadrat biomass from rake biomass. Overall relationship is shown in a) and effect of substrate and growth form are shown in b) and c) respectively. Regression line is represented by the dark solid line with 95% confidence intervals as light grey bands and error bars are standard errors. Quadrat and rake absence (value displayed at 0.1) are depicted in a) and b) but are not included in regressions calculation.

**Table 2:** Equations and summary statistics of models allowing to predict quadrat biomass (log_10_ g/m^2^) from rake biomass (log_10_ g/m^2^). Models are shown for all available data and for different subsets with distinct environmental conditions having a significant effect on the quadrat and rake relationship. Equ. equation number, RMSECV root mean square error of cross-validation, MAPE mean percentage absolute error.

| Equ. | Environmental condition | Coefficients | | RMSECV<br>($\log_{10}$, g/m<sup>2</sup>) | MAPE<br>(%) | R<sup>2</sup> | *n* | Rake biomass range<br>(g/m<sup>2</sup>) |
| --- | --- | --- | --- | --- | --- | --- | --- | --- |
|  |  | Intercept | Slope |  |  |  |  |  |
| 1 | All data | 0.91 | 0.68 | 0.34 | 21 | 0.62 | 67 | 0.36 - 1257.68 |
| 2 | Silt | 1.11 | 0.44 | 0.31 | 17 | 0.46 | 16 | 0.84 - 136.45 |
| 3 | Sand | 0.76 | 0.78 | 0.35 | 24 | 0.69 | 40 | 0.36 - 1257.68 |
| 4 | Rosette | 0.92 | 0.76 | 0.34 | 22 | 0.61 | 26 | 0.36 - 59.91 |
| 5 | Canopy | 0.63 |  |  |  |  | 20 | 2.30 - 1257.68 |

The discrepancy between rake and quadrat biomass comparison was not constant over the biomass gradient (Figure 3a, Table 2). The slope was less than one (0.68) and rake underestimation of quadrat biomass was highest at low biomass, while measurements from both methods converged at higher biomass (> ∼ 100 g/m^2^). The RMSECV of the prediction (2.19 g/m^2^) was similar to the standard error associated with the rake measurements and smaller than the error of the quadrat measurements. The MAPE of the predictions (21%) was lower than the relative error of both quadrat and rake biomass measurements (69% and 53% respectively). The intercept was greater than 0 (log_10_(quadrat biomass) = 0.91 or quadrat biomass = 8.13 g/m^2^), therefore application of the equation would result in systematically biased biomass estimation in the absence of SAV and has to be limited to presence data (Table 2). Nonetheless, the model yielded a good prediction of quadrat biomass with a similar mean and standard deviation compared to observations (Supplemental information - SI - Figure S1).

Predictions of quadrat biomass from rake measurements were affected by substrate type and SAV growth form (Figure 3b and 3c), while no effect of collection strategy, biomass treatment, lake or depth was detected. The slopes and intercepts of the relationship differed significantly among substrate type (*F*_2, 61_ = 4.44, *p* = 0.02), with silt displaying a higher intercept, lower slope and higher variability than sandy sediments (Table 1). This was indicative of the greater efficiency of rake to collect SAV, especially at low biomass, in sand bed areas compared to finer (silt) and more organic sediments (silt). For sandy sediments, the slope was not only closer to one, but the relationship also had a better fit. In hard packed sediments (clay - pebble), rake sampling seemed to be unsuited and completely failed to collect any plant despite their known presence. However, this substrate type occurred only at a reduced number of sites (*n* = 6) and the biomass at these sites was very low (< 0.5 g/m^2^).

In the case of the effect of the dominant growth form, the relationship slopes were similar for rosette and canopy-forming SAV (*F*_1,42_ = 0.02, *p* = 0.88), but their intercepts were significantly different (*F*_1,43_ = 6.93, *p* = 0.01). Understory were dominant at only 2 sites and were excluded from analysis. For the same rake biomass, the rosette had a quadrat biomass systematically higher than canopy by 1.95 g/m^2^. Thus, rake was more efficient at sampling canopy-forming plants and tended to underestimate the biomass of rosette-forming *Vallisneria americana*. Inclusion of the dominant SAV growth form in the model resulted in an overall better fit and lower prediction error.

### Determination of optimal spatial resolution between rake and echosounding

To determine the spatial resolution of rake biomass at which we could compare acoustic data, we inspected the relationships between rake biomass and mean biovolume at increasing radial distance from rake sites (Figure 4a). As expected, the correlation coefficients between biovolume and rake biomass were highest close to the rake site and decreased with distance. However, the difference in correlation with distance were subtle and the observed range narrow (*r*_min_ = 0.66, *r*_max_ = 0.74). Deviation from the general decreasing trend and slight increases in the strength of the correlation were observed starting at 7 m, with a maximum increase reached around 10 m. The correlation increased again around 25 m radius and decrease thereafter to reach a minimum correlation value. The overall decreasing correlations were computed using an increasing number of paired observations with distance, which also increased their significance level (Figure 4b). These results justified our selection of a 10-m radius for the spatial resolution of biovolume, corresponding to an intermediate correlation value (*r*_10m_ = 0.73) derived from a reasonably high number of paired observations (*n* = 52). Furthermore, a resolution area coinciding with a 10-m radius is consistent with field observations and rake site area, taking into consideration boat size (7 m long), its movements around the anchor, rake collection at 2 m from boat sides and positioning error (up to ∼ 3 m). The chosen echosounding resolution is hereafter referred to biovolume_10_.

**Figure 4:**
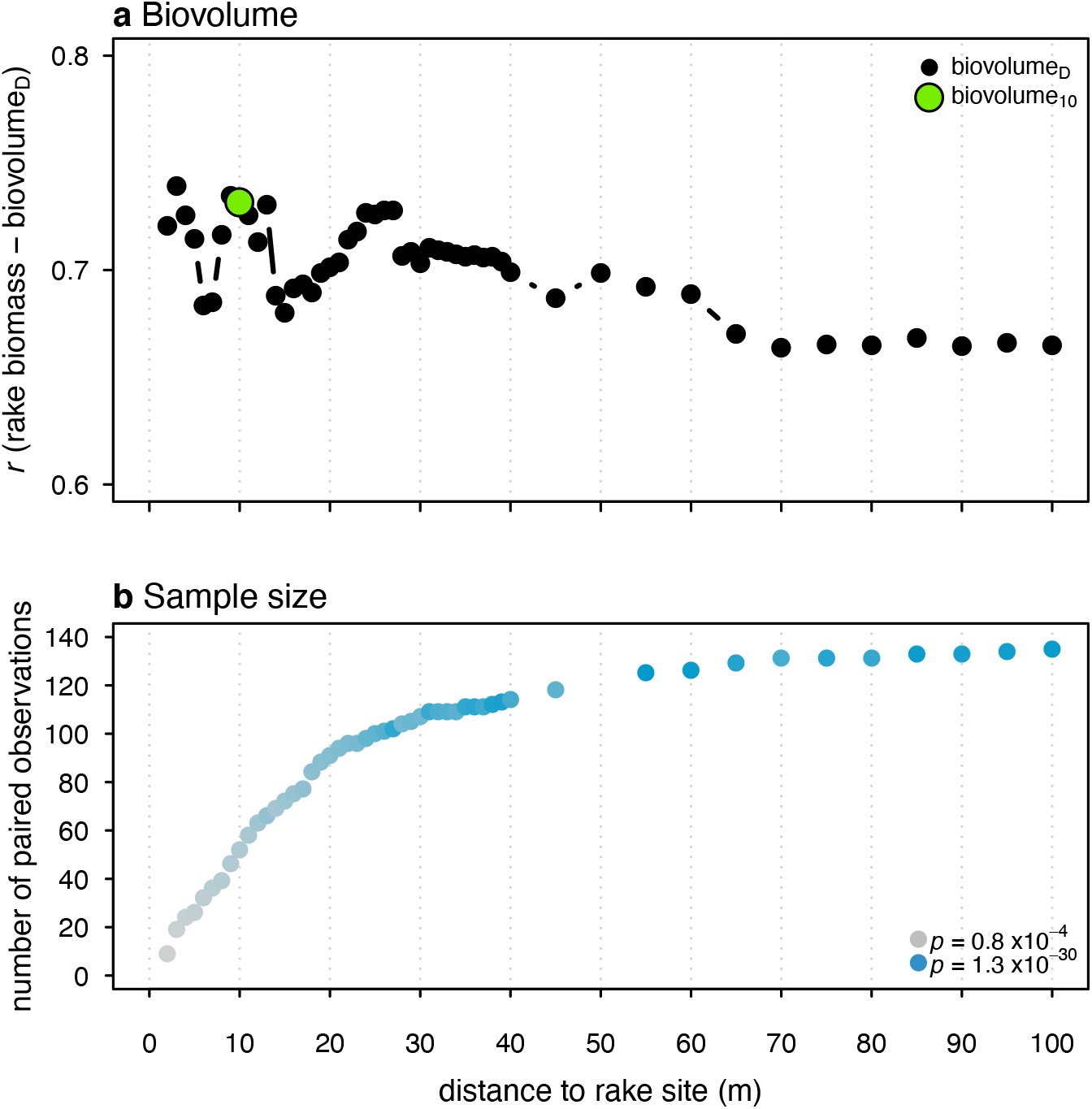
Effect of spatial resolution on the relationship strength between rake biomass and echosounding biomass proxy (biovolume). A) Pearson correlation (*r*) between rake biomass and mean biovolume aggregated at increasing radius distance from rake site (biovolume_D_). The green dot indicates the correlation at the selected resolution of 10-m radius. B) Number of paired observations and significance level for the correlations.

### Interannual variability of the paired echosounding and rake observations dataset

The resulting 52 pairs of acoustic and rake observations were collected during 7 independent field campaigns (spring 2012 and summers 2012-2017) that differed markedly in terms of SAV abundance and community composition due to interannual variation in environmental conditions (SI Figure S2). Of these 52 sites, 50 were used is subsequent analyses as 2 pairs were collected in depths too shallow for echosounding. This was found by using a 0.5 m threshold to depth collected during rake collection (Sabol et al. 2002); we chose rake depth since comparison to echosounding depth revealed the latter to be overestimated in shallow areas (<1.5 m, SI Figure S3). Only one site corresponded to a matched pair in 2016 and it was not included in the interannual statistical analysis. The rake biomass differed significantly among campaigns (*F*_5, 43_ = 3.78, *p* = 0.006, SI Figure S2a), with lowest biomass measured in spring 2012 and summer 2014 and maximum biomass measured in summers 2013 and 2015. Similar interannual patterns were observed for biovolume_10_ (*F*_5, 43_ = 6.59, *p* = 0.0001, SI Figure S2b). In contrast to SAV abundances, the growth form was generally equally distributed between canopy and rosette throughout the campaigns (*F*_5, 43_ = 2.25, *p* = 0.07, SI Figure S2c), with a notable exception in 2014 where rosette SAV (*Vallisneria americana*) was dominant. Understory (*Chara spp*) were present at only 3 sites at a relative abundance of less than 6% and were thus excluded from analysis. The filamentous algal mats biomasses were significantly different between campaigns (Chlorophytes *H* = 16.73, *p* = 0.005, *df* = 5; *Lyngbya H* = 14.33, *p* = 0.01), and were generally higher in years of high SAV biomass (2012, 2013, 2015, Figure SI S2d, e). Cyanobacterial mats (*Microseira wollei*) were also more abundant in 2017, coinciding with unusually high-water levels (x_level, echo2017_ = 1.22 m vs x_level, echo2012-2016_ = 0.43 m) as shown by the sharp differences in water depth recorded among campaigns (*F*_5, 43_ = 24.23, *p* < 0.0001, SI Figure S2f). Campaigns tended to be either characterized by deep (2014 and 2017), shallow (2012) or intermediate water depths (2013, 2015). Very high to intermediate water depths corresponded with abundant *Microseira* and an absence of Chlorophytes, while years (2013, 2015) of intermediate water depth coincided with high biomass of both *Microseira* and Chlorophytes. This is supported by previous reports of the impact of water level fluctuation on filamentous algal mat types, with lower level favouring Chlorophytes (Cattaneo et al. 2013) while *Microseira* prevails under low light intensity and high water levels (Cattaneo et al. 2013; Hudon et al. 2014).

We also investigated how acoustic data collection and quality varied among campaigns (SI Figure S4). Despite significant differences in wind speed among campaigns (*H* = 25.98, *p* < 0.001, *df* = 5, SI Figure S4a) reaching its highest values in spring 2012, the overall wind speed was generally low (*x̄* = 3.3 m/s) thus limiting the creation of turbulence-induced bubbles interfering with the SAV echo signal. As could be expected from dominant wind pattern, the wind direction was constant across campaigns, generally coming from the South-West (*H* = 2.75, *p* = 0.73, *df* = 5), with somewhat more variability in 2014 and 2017 (SI Figure S4b). Although wind conditions were favorable for data collection, the acoustic data quality was variable among campaigns. The error associated with biovolume_10_ estimation, i.e. the biovolume standard deviation (biovolume_10_ sd), was stable between years (*H* = 4.06, *p* = 0.54, *df* = 5) but had a lower variance in 2012 than over the other years (SI Figure S4c). This could result from the higher sampling effort and higher number of ping reports per site in 2012, which generally decreased over subsequent years, inducing significantly lower sampling frequencies between campaigns (*H* = 21.61, *p* = 0.0001, *df* = 5, SI Figure S4d). The number of ping reports was generally 7 times higher in June 2012 than during the lower sampling effort campaign of 2017 (median_J2012_ = 71, median_2017_ = 10). This reduced frequency was accompanied by a reduced proximity between rake and acoustic data across campaigns, although there were no significant differences in proximity (*F*_5, 43_ = 0.70, *p* = 0.63, SI Figure S4e). The echosounding survey tracks were typically going back and forth close to the rake site in 2012 and 2013 and were thus closer with medians of 6 m from the rake sites, compared to 9 m in 2017. The more precise positioning in earlier years explains the higher number of paired rake-acoustic observations (*n* = 17 to 10 for 2012 to 2014) compared to more recent years (*n* = 6 and 4 for 2015 and 2017).

### Prediction of rake biomass from acoustic data

To predict rake biomass from biovolume_10_ and test for the influence of various environmental conditions on the relationship, we performed a PLSR. We first selected the number of components (or latent variables) using RMSECV which had a minimal value of 0.60 (log_10_) at 8 components (Figure 5a). To avoid overfitting, we chose the model with three components (RMSECV = 0.63), which was the model with the fewest components that was less than one standard deviation from the best model (one-sigma heuristic approach). Biovolume_10_, biovolume_10_ sd and canopy growth form were the most important variables explaining these components (Figure 5b). When looking at site configuration in relation to predictors, biovolume_10_ and biovolume_10_ sd were nearly orthogonal (Figure 5c). Biovolume was more highly correlated to rake SAV biomass and canopy growth form, while biovolume_10_ sd was pulled by two extreme data points. The two echosounding variables were correlated to rake biomass (*r*_biovolume_ = 0.79, *r*_biovolume sd_ = 0.61, SI Figure S5a, b) but also to one another (*r* = 0.59, SI Figure S5c). Above a biovolume of 15, there was increased scatter between the two variables and a reduced correlation (*r*_biovolume > 4_ = 0.10). The presence of canopy growth form modified the relationships between these predictors and rake biomass, where canopy had a higher rake biomass per biovolume_10_ than rosette-forming plants (SI Figure S5d). Wind and echosounding data quality variables also significantly discriminated between components (*p* < 0.05, Figure 5b), but they had lower SR and their effect on the relationship between biovolume_10_ and rake biomass was evident only for depth and wind direction. Shallow sites (< 1 m) tended to display a low biovolume^10^ for a given rake biomass, while sites sampled when winds were coming from North to North-West had high biovolume_10_ for their measured rake biomass (Figure S5e, f).

**Figure 5:**
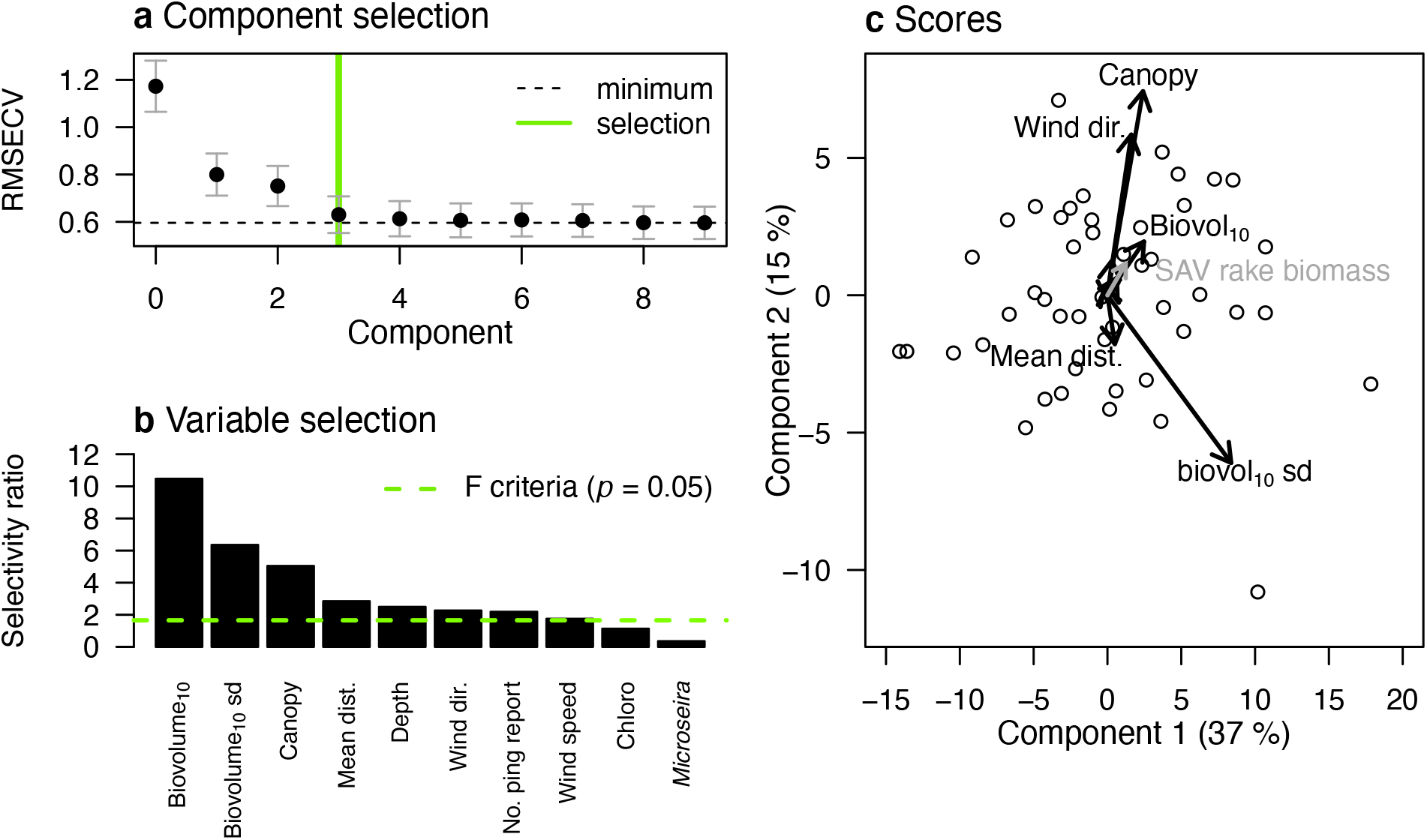
Component and variable selection for the PLSR model including all possible predictors of rake biomass. A) Component selection using RMSECV. B) Predictors importance for three components using selectivity ratio. C) Score plot describing data configuration in relation to predictors and with the explained variable in grey. Error bars are the standard error of the residuals and numbers in parentheses are the variance explained by the component. In b, bars above the dashed line have a significant discriminant power at *p* < 0.05. RMSECV root mean square error of cross validation, sd standard deviation, no. number, dist. distance, dir. direction.

Components were a decomposition of the original variables and the highest quality of predictions were reached at 3 components. Therefore, to build our prediction models, we chose the three most important variables explaining these components and that were also easily measured in practice. We created two models, with or without proportion of canopy-forming SAV, since species identification is not always readily available at large spatial scales. The final models with the selected variables yielded the following equations:

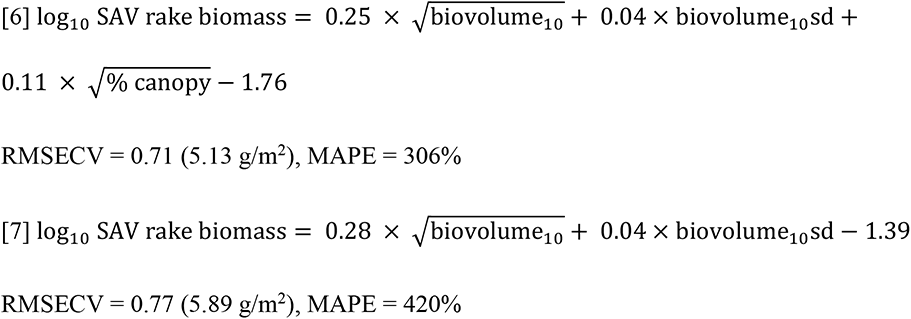

The relationship between rake biomass and biovolume_10_ was moderately strong with an R^2^ of 0.53 or 0.66 when including SAV growth form. The absolute error (RMSECV) of the models was similar to the error of the rake collection (*x̄*_SE,rake_ = 6.82 g/m^2^). However, the predictions had a higher relative error (MAPE) than the relative error of both the rake biomass and biovolume_10_, which was relatively high given the low measurement range (83 %, rake biomass range 0.002 – 125.50 g/m^2^, 86% biovolume_10_ range 0.19 – 109 m). SAV appeared to be more spatially aggregated when looking at biovolume_10_ with 98% of observations having a variance higher than its mean (s^2^ > *x̄*) compared to 50% for rake. Nonetheless, there was no systematic bias of the predictions, exactly half of the predictions were either above or below its paired observation when using the model with 2 variables only (Figure S6). This yielded an overall good prediction of our sample, with a similar distribution between observations and predictions. Although the difference was not significant, the predictions had a lower mean and a higher standard deviation than the observations (*x̄*_obs_ = 17.33 g/m^2^, sd_obs_ = 29.92 g/m^2^, *x̄*_pred_ = 20.90 g/m^2^, sd_pred_ = 68,15 g/m^2^).

### Two-step model validation

Predicted quadrat biomass was determined by sequentially applying the simple two variables acoustic to rake biomass equation followed by the rake-quadrat equation. Predicted biomass was then compared to measured quadrat biomass. Three surveys carried out over different depth ranges were available for the comparison: a small-scale survey at constant depth (2013, 1.4 - 1.5 m), a survey in the deeper water in the North of LSP (0.5 - 7.0 m) and a survey in the shallow waters in the South of LSP (< 2.7 m). We first compared paired acoustic predictions at 10 m distance of quadrat measurements (Figure 6a). The two step predictions were comparable to the quadrat measurement and did not introduce any evident bias. The standard deviation of the Monte Carlo simulation was 3.89 g/m^2^ (log_10_ 0.59) which was an intermediate value between the prediction error of the two models. However, the RMSE at 122.65 g/m^2^ was higher than the quadrat measured standard error (38.02 g/m^2^). The MAPE of the model was also higher than the relative error of the quadrat measurements (1718% vs 50% respectively), but it was driven by a single outlying data point in the North of LSP that, once removed yielded a MAPE of 87%. Indeed, when looking at each individual survey, the acoustic predictions in 2016 tended to have a higher error than the 2013 predictions (RMSE_2016_ = 170.23 g/m^2^, MAPE_2016_ = 6433 %, RMSE_2013_ = 85.89 g/m^2^, MAPE_2013_ = 68 %). The 2016 predictions were either well above or below the 1:1 line, while the 2013 predictions from sites at a constant depth had a majority of their confidence intervals overlapping the 1:1 line. In 2016, sites from the deeper North sector tended to have higher predictions compared to nearly absent quadrat biomass (< 1 g/m^2^). Echosounding integrates a larger number of measured units and thus better describes the same areal extent than quadrat (up to 400 m^2^ vs < 1 m^2^) which, in this case, probably underestimates true biomass. Conversely, sites from the shallow South sector tended to have much lower acoustic predictions than that measured by quadrat, probably due to a bias from echosounding due to biomass accumulation at the surface from floating leaves. Overall, the predictions were not significantly different from quadrat measurements (paired t-test *p* = 0.17, Figure 6b), although acoustic predictions had somewhat lower mean and median (*x̄*_pred_ = 88.16 g/m^2^, *x̄*_obs_ = 121.40 g/m^2^, median_pred_ = 98.06 g/m2, median_obs_ = 107.31 g/m^2^).

**Figure 6:**
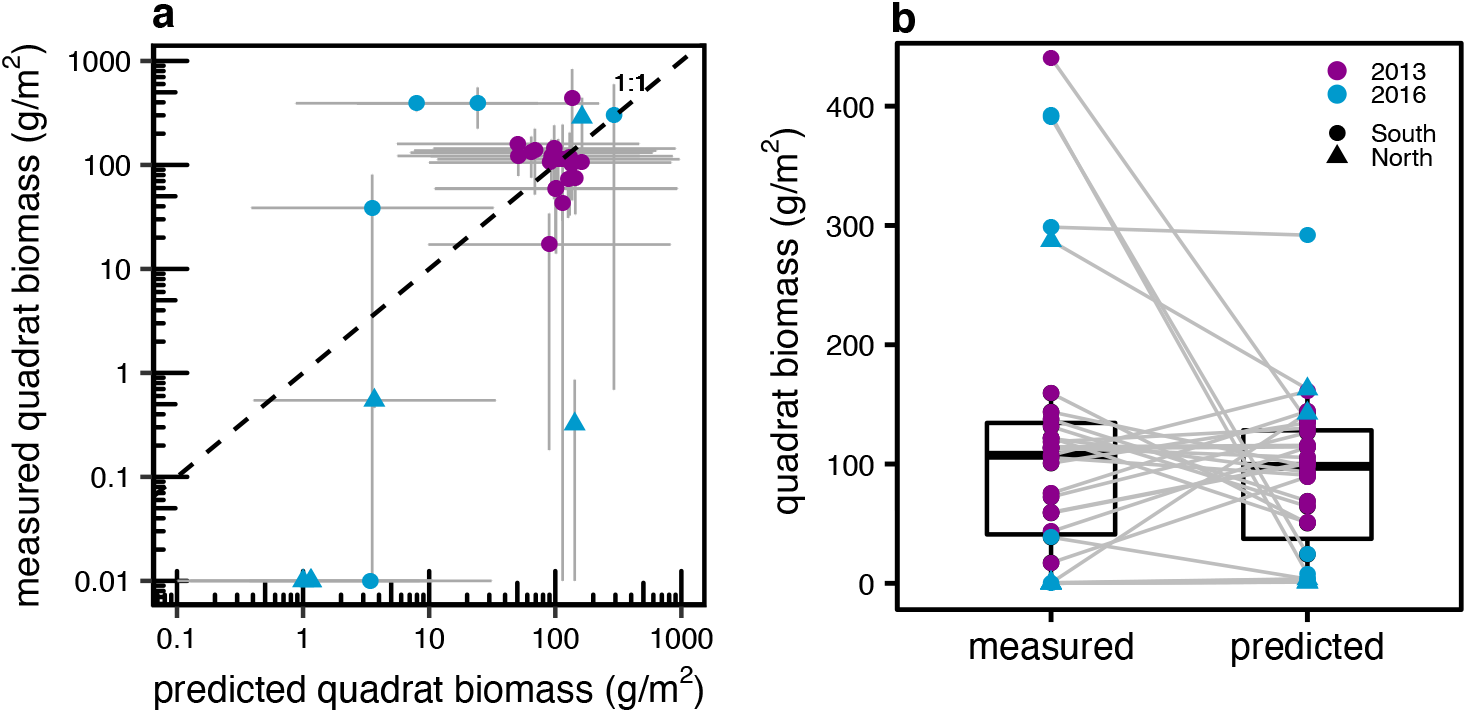
Validation of the two-step quadrat biomass prediction from echosounding (using equation 1, table 2 and equation 6 in text). A) Scatterplot of measured vs predicted quadrat biomasses, error bars are the 95% confidence intervals and absence of measured biomass are depicted at 0.01 g/m^2^. B) Paired boxplot of measured vs predicted quadrat biomasses, solid horizontal line within boxes represents the median, boxes extent the 25th and 75th percentiles and whiskers the 10th and 90th percentiles.

Second, given that most quadrat measurements in our dataset were located beyond 10 m from the echosounding track, we compared spatially interpolated measured quadrat biomass to interpolated predictions derived from echosounding (Figure 7). In the 2013 constant depth range survey, the quadrat and echosounding estimation were very similar and small local differences were potentially caused by the higher sampling effort and surveyed area of echosounding (Figure 7a, b). In the deeper North sector of LSP, interpolated biomasses were similar for quadrat and echosounding above the SAV maximum colonization depth (3 m, Figure 7c, d). Echosounding clearly enabled the delimitation of SAV spatially because of a better accountability of limits imposed by depths. Quadrat interpolation tended to overestimate biomass in deeper waters where no sampling occurred. Conversely, in the shallow South (< 1.5 m, Figure 7e, f), echosounding underestimated biomass, again probably because of bent SAV that accumulated at the water surface and prevented efficient estimation of underwater biomass using acoustic signal.

**Figure 7:**
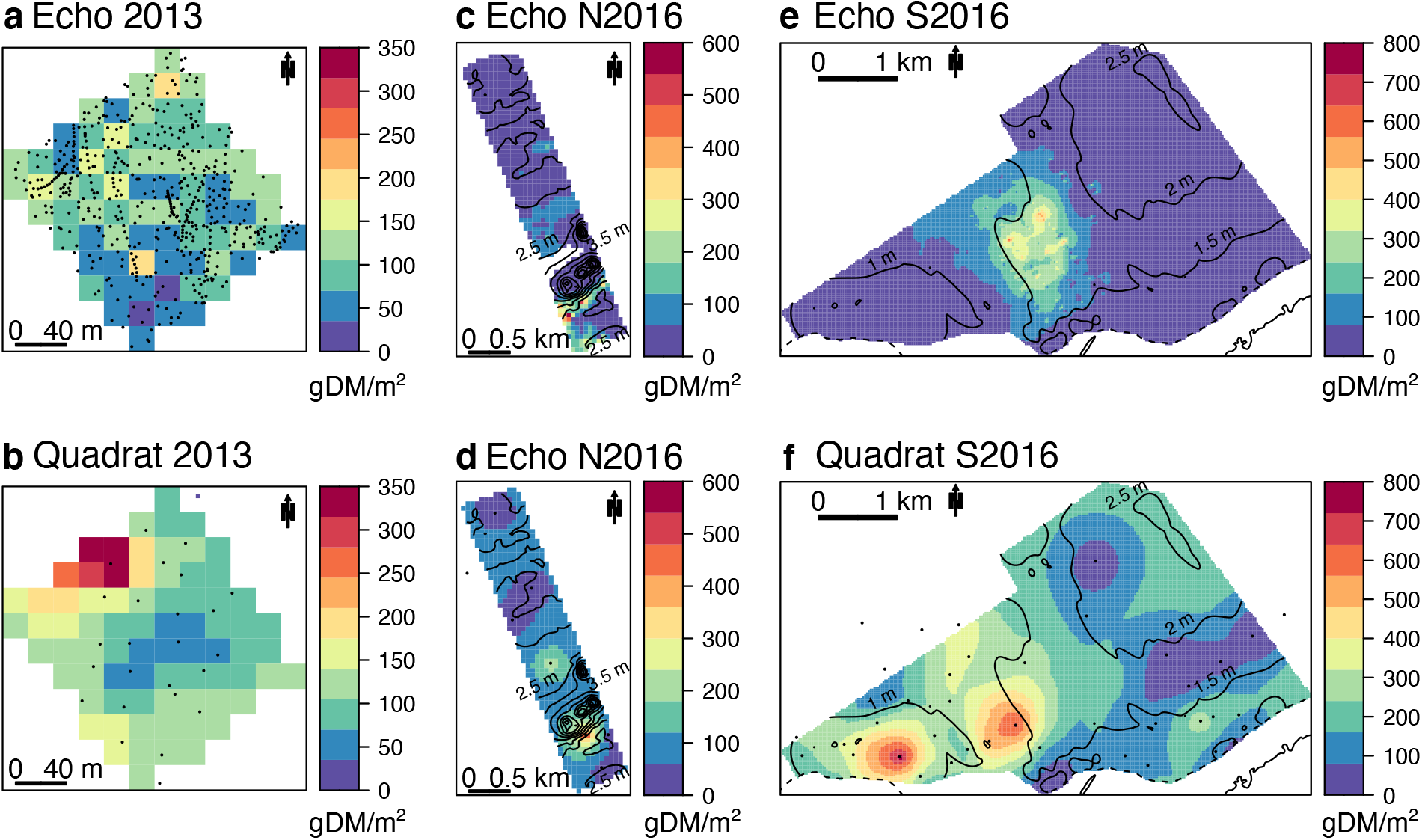
Comparison of spatially interpolated quadrat biomass estimated from echosounding (top panels) and point measurements (bottom panels). A and b 2013 small-scale plot of constant depth, c and d North section of LSP with depths ranging from 0.5 to 7 m sampled in 2016, e and f South section of LSP with shallow depths (< 2.5 m). Lines are isobaths at 0.5 m increments and dots sampled sites (not displayed for clarity in c and e). DM dry mass.

### Effect of sample size on whole system estimation

Finally, we assessed how the increased sample size afforded by echosounding and the use of spatial interpolation method modify whole-system biomass estimation. Using the two models we developed, we predicted quadrat biomass from both rake biomass and biovolume_10_ for five independent SAV surveys where both rake sampling and echosounding had similar spatial extent. We then compared the mean from these estimated biomasses, either on the raw data or on spatially interpolated biomass (Figure 8). In all surveyed years, the study area displayed heterogenous biomasses, with clear high and low biomass zones (SI Figure S7). This combined with the different sampling effort of rake (n = 21 to 30) and echosounding (n = 1224 to 1934) created widely different mean biomass per survey. Biomass predicted from rake was, depending on the survey, either lower or higher than that from echosounding generally by a factor of 2. The mean estimate of each technique was distinct and there was almost no overlap with their confidence intervals. As a result, the range across years of mean biomass from rake (14.61 to 59.20 g/m^2^) was more limited compared to that from echosounding (8.67 to 83.97 g/m^2^). The higher number of observations using echosounding generated more precise estimates with smaller confidence intervals compared to the very large uncertainty associated with the rake estimation. In contrast to the difference between techniques, spatial interpolation of biomass derived from echosounding did not affect whole-system mean biomass that was very similar to direct estimation from the original echosounding track. Largest differences between estimates were observed for surveys with interrupted echosounding tracks (2013, 2017) that consequently were not covering uniformly the study area (SI Figure S8).

**Figure 8:**
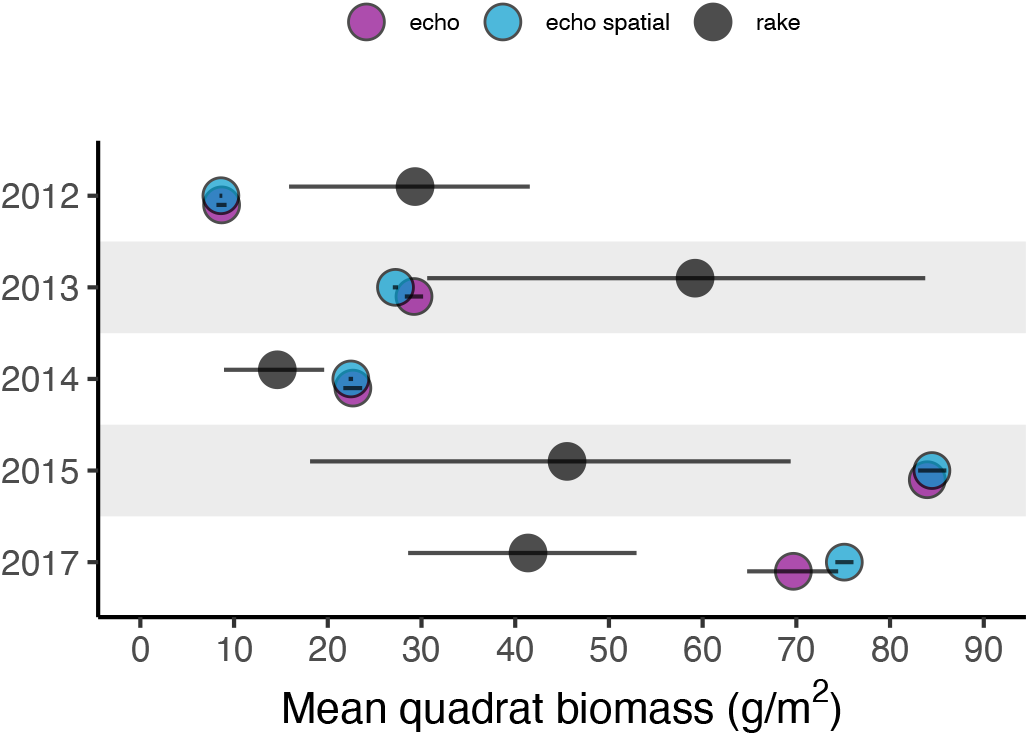
Comparison of whole-system average quadrat biomass using non-spatial predictions from rake and echosounding and spatially interpolated echosounding predicted biomass for five summertime yearly surveys. Horizontal bars represent 95% bootstrapped confidence intervals.

## Discussion

We successfully developed two intercalibrations that allows for the interchangeable use of quadrat, rake and echosounding techniques to estimate SAV biomass. We first predicted quadrat biomass from rake biomass and showed the effect of substrate and species growth form on the predictions. Using a resolution of 10 m radius, we predicted rake biomass from biovolume, a proxy of biomass derived from echosounding. We also showed that this prediction is affected by SAV growth form, depth, wind conditions and acoustic data quality. By sequentially applying both models to echosounding tracks, we were able to accurately predict quadrat biomass. By correcting its bias, rake thus has the potential to be a versatile ground truthing techniques for echosounding. In rugged and deeper bottoms, echosounding outperformed point sampling techniques in estimating biomass, but underestimated biomass in very shallow waters. Use of echosounding combined with the intercalibrations are particularly useful at large spatial scales as the higher sampling effort from the greater number of observations provided by this technology increase accuracy by capturing SAV heterogeneity.

### Intercalibration between quadrat and rake

Our intercalibration between quadrat and rake biomass confirmed that the two techniques are comparable, and that bias introduced by rake sampling can be corrected (Johnson and Newman 2011; Kenow et al. 2007; Rodusky et al. 2005). This correction is important and meaningful, since failure to correct biomass estimation from the rake can lead to a 4-fold underestimation of biomass, with even greater bias at low biomass. In real-world application, the correction can modify sample distribution which can reveal significant difference or patterns in space and times that would not be detected otherwise. Given that the correction is stronger below a rake biomass of 100 g/m^2^, these effects will depend on the measured rake biomass ranges. The model error was also lower than the standard error of the measured quadrat biomass and was equivalent to the error associated with rake biomass collection. The smaller error and variance of rake were probably caused by the larger rake sample unit size that dampened small-scale heterogeneity captured by quadrat, which had consistently higher ratio of variance to mean biomass. Thus, gain in accuracy, did not come at the expense of precision.

Our founding that rake collection underestimated biomass confirms previous observations on rake and quadrat comparisons (Johnson and Newman 2011; Kenow et al. 2007; Rodusky et al. 2005). This bias is probably introduced by saturation of plant material on the rake or to the loss of this material as it is lifted from the water. Work by Masto et al. (2020), who combined quadrat-rake apparatus and picked up remaining plant material after rake collection, also suggests that the rake does not completely break plant material at the sediment surface. This harvesting efficiency from rake was affected by the same factors explaining SAV anchorage strength from the natural pulling forces of waves, current or bird foraging: the size of SAV root system and sediment cohesive strength (Schutten et al. 2005). Indeed, we found that canopy-forming SAV tended to be more efficiently collected by rake in contrast to the rosette-forming *Vallisneria americana*. The latter has one of the higher root to shoot ratio among freshwater SAV (Stevenson 1988), thus being harder to break from the sediment and being systematically underestimated by the rake. We also observed that the rosette-forming linear leaves tended slip in between rake teeth compared to the canopy-forming that were entangled in them. Additionally, canopy-forming not only tend to have a reduced root system, but their intertwined stem could drag plant material from outside the rake sampled area. Dense stands of the canopy-forming species *Ceratophyllum demersum*, *Potamogeton zosteriformi*, *Hydrilla verticillata* have previously been overestimated by rake techniques (Johnson and Newman 2011; Rodusky et al. 2005).

Our results further indicate that rake harvesting efficiency is dependent on the substrate type. We did find that in hard packed sediments (pebble-clay), which provide higher anchorage strength, the rake technique failed to collect any SAV. Counterintuitively, plants were more easily and consistently pulled out from moderately compacted (sand) than from organic and soft sediments (silt). This finding could be an artifact of the more restricted biomass range measured from siltier sites as compared to sandier ones in our survey. However, Rodusky et al. (2005) similarly found a weaker relationship with quadrat biomass and a lower slope when rake collection was on peat-like organic sediments compared to sand. In very loose and organic sediments, the rake could have less grip and SAV be more elusive, being both dragged and buried in mud by the rake motion. SAV is also likely more dispersed in this type of substrate since it is not optimal for their growth (Barko et al. 1991). All of these effects would result in less consistent rake harvest in organic substrates. Therefore, calibration of rake biomass measurements is site specific and inclusion of both species and substrate information increase accuracy. Nevertheless, we have shown that the calibration is not impacted by the sampling strategy and the depth of the quadrat and rake comparison. Investigators can thus use the simplest sampling strategy, such as the haphazard pair, and reduce sampling effort in deeper more hazardous areas for scuba diving.

### Intercalibration between rake and echosounding

We also successfully developed an intercalibration between rake biomass and biovolume_10_. Quadrat biomass has previously been related to biovolume or height measured by echosounder (e.g. Duarte 1987; Maceina et al. 1984; Thomas et al. 1990), but to our knowledge, only Howell and Richardson (2019) related rake biomass to biovolume. However, their biovolume was derived from a cloud-based data-processing platform and was not an absolute volumetric estimate; rather it referred to the percent volume inhabited by SAV in the water column (PVI, % cover/depth * height, (Thomas et al. 1990; Winfield et al. 2007). This metric is not appropriate to estimate biomass because it is a depth-standardized measurement that does not reflect variation in plant height. For example, given a surface cover of 100%, the same PVI of 80% is measured for a plant height of 2.4 m at a depth of 3 m (2.4/3 * 100) or a height of 0.8 m at a depth of 1 m (0.8/1 * 100). Comparison of these specimens’ biomass would likely be very different given their 3-fold size difference. Thus, care must be taken when blindly applying measure from the manufacturer; the use of the simple biovolume as % cover multiplied by height is recommended to compare with biomass estimates.

Our comparison yielded a simple model with three predictors of biomass: biovolume_10_, its associated error and the proportion of canopy-forming SAV. Indeed, Duarte (1987) has previously shown that given their different biomass allocation with height, deriving calibration by SAV growth form increase accuracy in quadrat biomass estimation. Although not detected in our model, filamentous algae could further introduce bias in the biovolume-rake biomass relationship since they can be detected by echosounder (Bučas et al. 2016; Depew et al. 2009). Predictions were also affected by environmental conditions during sampling, with clear effect of depth and wind direction. The depth effect could be due to differences in species composition that tend to vary by depth at our study site (Hudon 1997). Alternatively, underestimation in shallow areas could be due to detection problems either because of an overestimation of bottom depth, which is used to calculate plant height, or of floating plant material at the water surface that was not detected by the echosounder. Observations when wind was coming from North to North-West also had overestimated biovolume_10_. In LSP, wind coming from this direction are against water current and are in the general orientation of the lake, which increases the travelling distance on open water. Both of these factors can increase wave height and create bubbles that cause false echosounding plant detection. To account for these environmental variations, greater accuracy could thus be provided by conducting a calibration per sampling campaign, for similar environmental conditions or during favorable wind conditions.

### Versatility of the approach

Our two step intercalibration model approach has several advantages over using techniques in isolation. First, it allows using rake to report accurate quadrat-equivalent biomass. This saves time since that, depending on the diver’s experience, a single quadrat collection can take up to half an hour (Downing and Anderson 1985) compared to a few minutes for rake. Furthermore, calibration between rake and quadrat can become a one-time event that prevents the need for scuba training and repeated hazardous sampling. Time and resources gained by using the rake in combination with the improved underwater accessibility can be dedicated to additional sample collection, thus improving sample size, understanding of SAV diversity, and spatial coverage.

Second, the two-step model approach allows for the interchangeable use of rake and echosounding. Comparing rake with echosounding is a challenge because the two techniques have very different number of samples collected over a given area. Therefore, we provide for the first time a resolution at which echosounding can be compared to rake sampling: 10 m radius (20 m diameter). The resolution from this remote sensing technique is similar to that of different satellite products, such as NASA Landsat (30 m) or ESA Sentinel-2 (10 m). However, this determined resolution is dependent on the rake sampling and could be improved. With more precise positioning and increased spatial proximity between rake replicates, resolution can be decreased or even chosen by the user. Furthermore, the calibration allows using the techniques where they perform best: in very shallow area for rake and in deeper areas for echosounding.

Third, by showing the association between echosounding and rake, and correcting rake bias, the intercalibrations make rake an appropriate and rapid ground truthing technique for echosounding. Indeed, the relationship between biovolume and biomass is species- or stand-specific and needs to be repeated with any changes in SAV species composition, including filamentous algae. This new ground truthing provides the opportunity to use echosounding to cover SAV meadows in large extents in a uniform manner, thus capturing more of the spatial heterogeneity and increasing accuracy at the ecosystem scale. In our whole-system estimation, this yielded increased ranges in biomass and differences among years. Although the precision of biomass predictions from echosounding at an individual site is lower than that of rake, that loss is counteracted by the sheer number of echosounding measurements. Reaching similar sampling effort would be impossible with the rake technique. We estimated that it would take 60 days using a rake to simply collect a similar ecosystem-scale sample size as observed in Figure 8, without taking into account processing time. We also saw that there is no real gain in using spatial interpolation techniques to estimate biomass if there is a thorough coverage of the study area with echosounding. This is more effective as it requires less post-processing and computing power. Furthermore, using the rake in combination with echosounding provides information on SAV species that is not available using echosounding.

## Comments and recommendations

The choice to use one or both intercalibrations will depend on the study goals, resources available and time granted. If one only intends to use rake, the quadrat and rake calibration can be conducted using the simplest sampling strategy, where only a one-time calibration is needed, with sites chosen to maximize a biomass gradient and ensure divers security. Increased accuracy can be achieved by deriving species- and substrate-specific equations. For the rake and echosounding combination, an even more efficient intercalibration can be achieved than the one presented here. For example, despite having a rake dataset of 217 sites, we only had 52 sites that spatially matched the echosounding tracks at 10 m radius resolution. Since the two sampling techniques were not used simultaneously, sampling with both techniques had to be conducted as close as possible to an assigned geographic position. Over the 6-year sampling period, variations in staff and GPS accuracy resulted in reduced precision, yielding a lower number of matched pairs in later years, which affected predictions (Figure 5b). These shortcomings can be overcome by conducting the rake sampling and echosounding simultaneously. For example, buoys could be deployed as the echosounder is passing over the selected calibration sites for subsequent rake collection. Furthermore, when dense plant material is floating at the water surface, echosounding is inappropriate: the transducer is blinded which leads to drastically underestimated biomass. Therefore, to maximize matched rake-echosounding sites and avoid sampling in inappropriate conditions, careful planning of sampling and training of field technicians are necessary.

Information needed regarding species composition, substrate type and time dedicated for biomass samples or echosounding post-processing should also be planned in advance. The rake has the advantage of providing species information, but each sample entails considerable sample processing time. Using a faster semi-quantitative approach such as applying a visual abundance scale based on the degree of each species filling the rake teeth can help increase sample size and provide community information at the ecosystem scale (Yin and Kreiling 2011), which can be used to more accurately predict biomass. When different SAV communities display distinctive heights, for example understory charophytes and taller angiosperms, applying height threshold to echosounding data can provide functional group information at large spatial scales (Bučas et al. 2016). Recent echosounders also provide information on substrate type that can increase accuracy when applying our two-step calibration approach (Helminen et al. 2019; Munday et al. 2013). To reduce purchase cost and post-processing time of echosounding, consumer-grade echosounder coupled to cloud-based automated data-processing tools can also be used (Helminen et al. 2019; Howell and Richardson 2019; Munday et al. 2013). New technological development in hydroacoustic autonomous boats (e.g. Goulon et al. 2021) are further promising to reduce sampling time and increase sampling frequency. Given the technological progress that were accomplished over the past decade, it is likely that future developments in GPS, echosounding and computing power will further facilitate and decrease costs related to large-scale estimations of SAV biomass over a wide range of environmental conditions.

## Acknowledgements

We thank Stéphanie Massé, Caroline Chartier, Elena Neszvecsko, Geneviève Trottier, Magalie Noiseux-Laurin and François Messier from GRIL, Patricia Bolduc from UQTR and all other students that conducted field work and processed samples. The technical support of Jean-Pierre Amyot, Conrad Beauvais, Rollando Benjamin, Maude Lachapelle, Yasmina Remmal and Maxime Wauthy from Environment Canada is acknowledged with thanks. Bruce Gray and the Technical Operations diving team (Environment Canada, Canada Centre for Inland Waters, Burlington, ON) provided excellent field support for underwater sample collection. We also thank Richard Carignan and Andrea Bertolo for planning and troubleshooting during echosounding surveys. Sampling was made possible thanks to the guides from pourvoirie Roger Gladu. This project was funded by the GRIL, a FRQNT Initiatives stratégiques pour l’innovation grant, FRQNT and NSERC scholarships to MB. This work was supported by Environment and Climate Change Canada (ECCC) as part of the Canada-Québec St. Lawrence Action Plan (SLAP) (CH).

---

**Figure S1:**
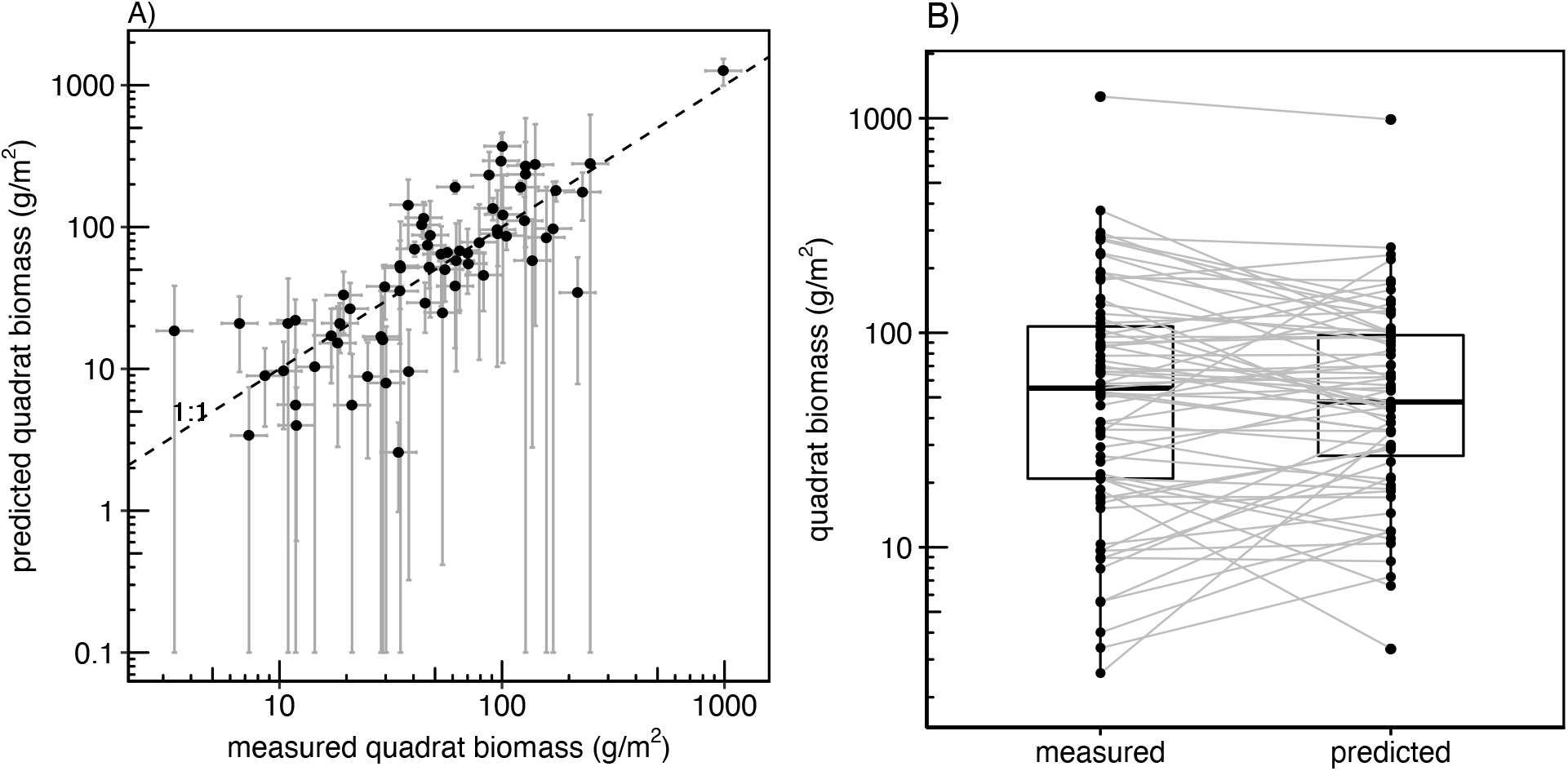
Evaluation of the cross-validation prediction of quadrat biomass from rake biomass. A) Scatterplot of measured vs predicted quadrat biomass, error bars are the 95% confidence interval. Lower limit of prediction interval below zero are depicted 0.1. B) Paired boxplot of measured vs predicted quadrat biomass, solid horizontal line within boxes represents the median, boxes extent the 25^th^ and 75^th^ percentiles and whiskers the 10^th^ and 90^th^ percentiles.

**Figure S2:**
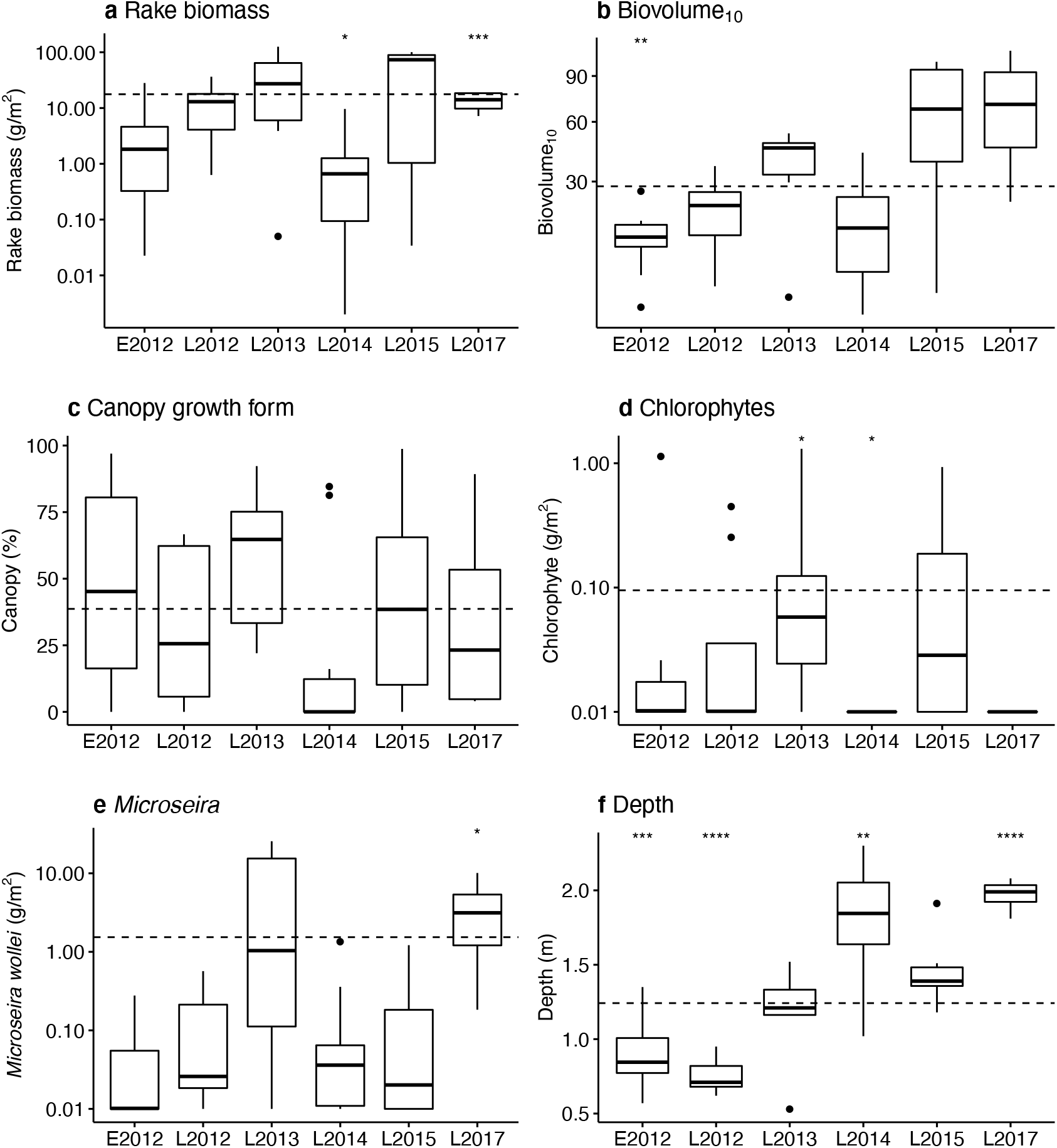
Boxplot of the E-R dataset variables per sampling campaign. E stand for early (June) and L for late summer campaign (July, August). The solid horizontal line within boxes represents the median, boxes extent the 25th and 75th percentiles and whiskers the 10th and 90th percentiles. The dashed line is the overall mean. Symbols represent significance levels of T tests or Wilcoxon tests between each group and the overall mean of the other groups at * *p* < 0.5, ** *p* < 0.01, *** *p* < 0.001, **** *p* < 0.0001. In d and f, absent filamentous algae biomass is depicted at 0.01 g/m^2^.

**Figure S3:**
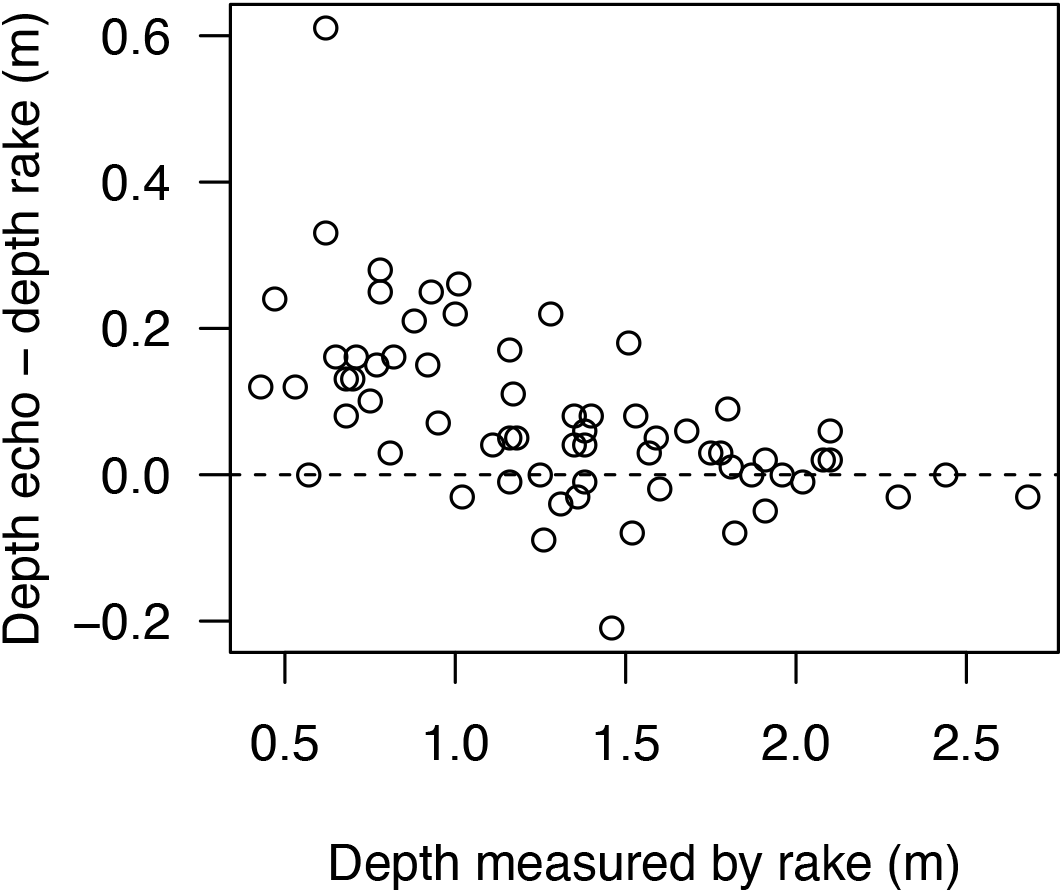
Comparison of depth measured by echosounding (echo) to depth measured during rake sampling. The latter is reported for water level at echosounding date.

**Figure S4:**
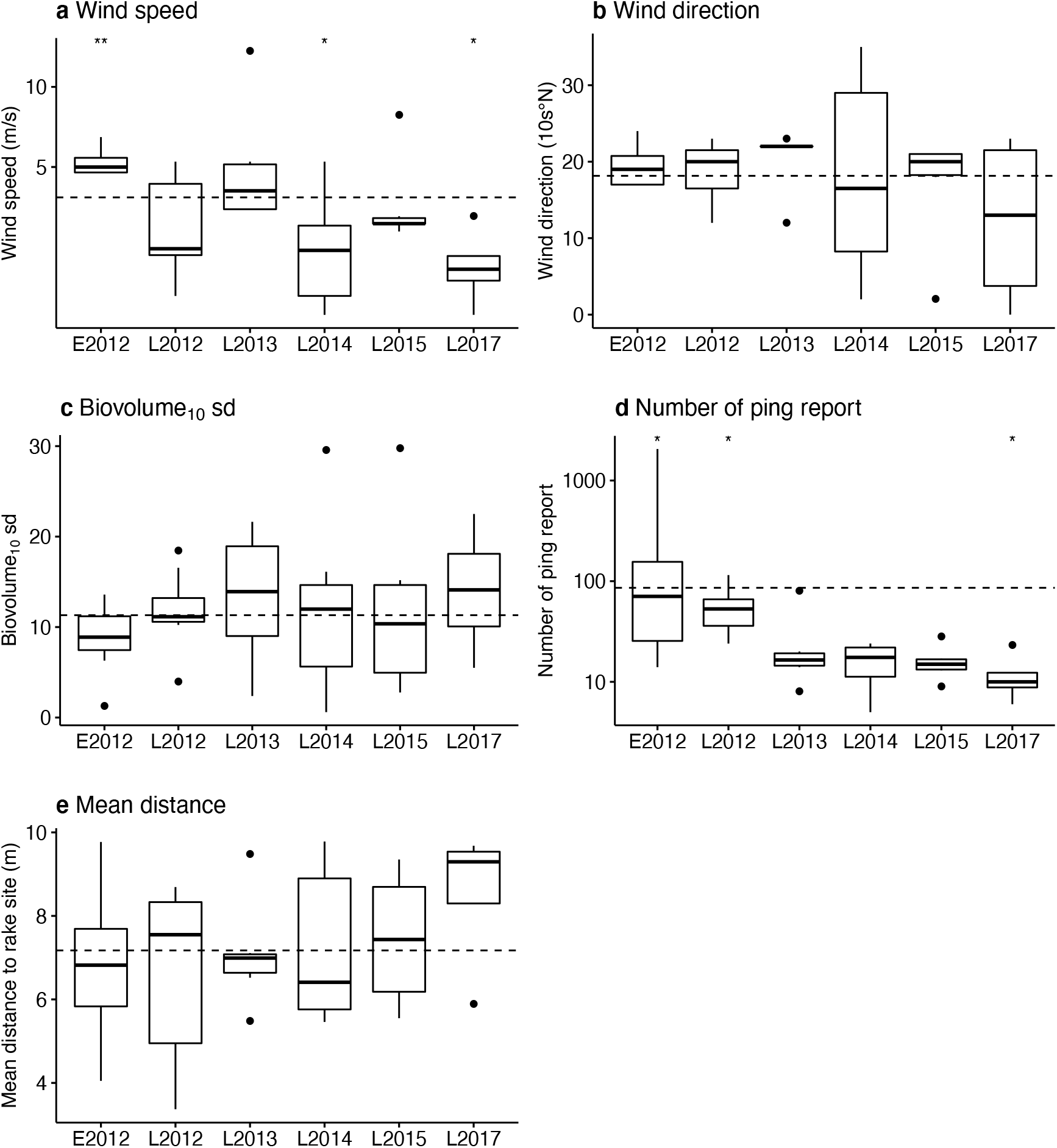
Boxplot of the E-R dataset variables per sampling campaign. S stand for summer campaign (July, August) and E early (June). The solid horizontal line within boxes represents the median, boxes extent the 25th and 75th percentiles and whiskers the 10th and 90th percentiles. The dashed line is the overall mean. Symbols represent significance levels of T tests or Wilcoxon tests between each group and the overall mean of the other groups at * *p* < 0.5, ** *p* < 0.01.

**Figure S5:**
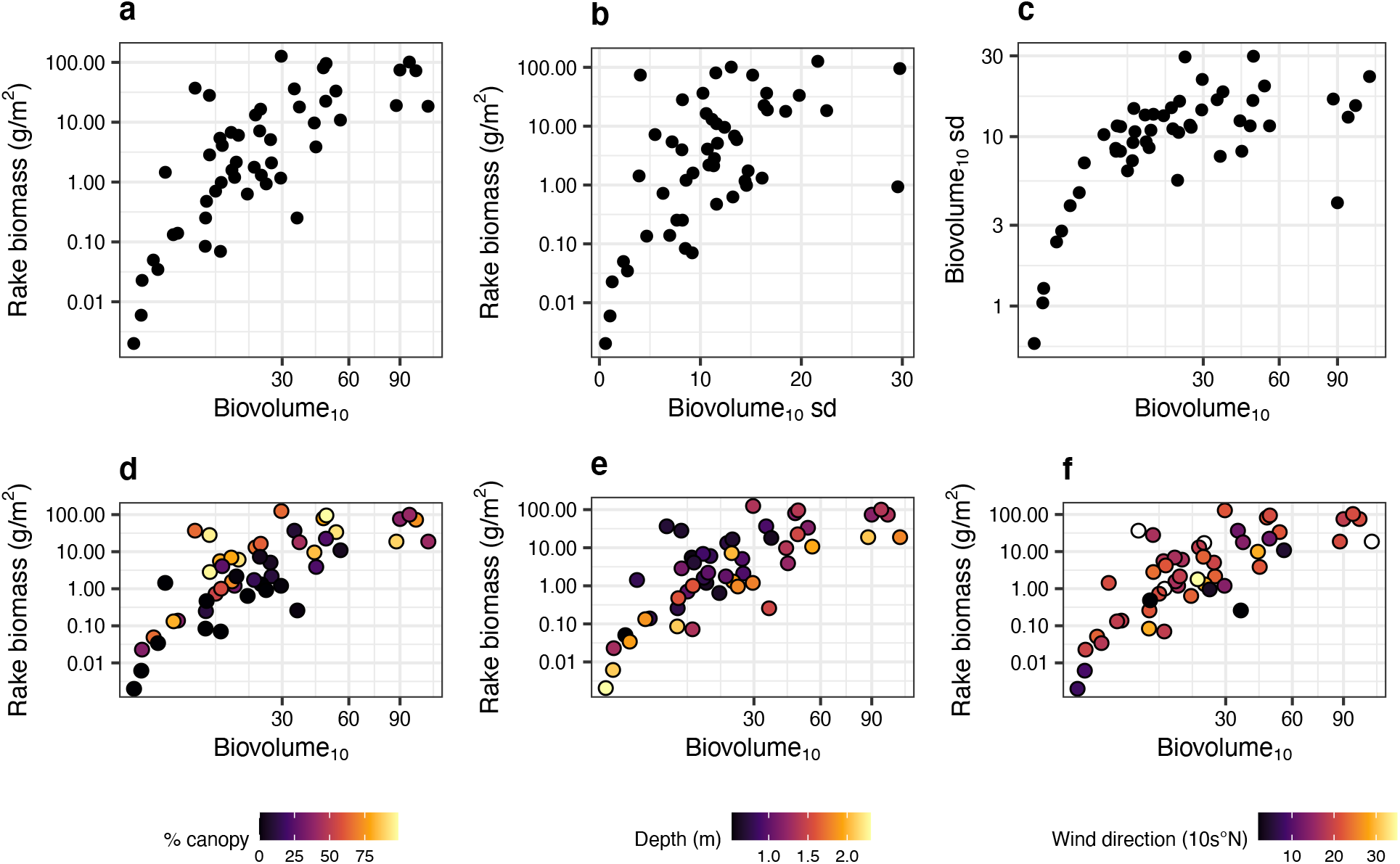
Scatterplot of the variables with higher discriminant power in the PLSR model. Biovolume_10_ (a) and biovolume_10_ sd (b) are predictors of rake biomass and are correlated to one another (c). Other variables (d - % canopy, e- depth, f- wind direction) modify the biovolume-rake biomass relationships. In f, Est is 9, South is 18, West is 27, North is 36 and 0, which indicate calm wind conditions, is not colored (n = 1).

**Figure S6:**
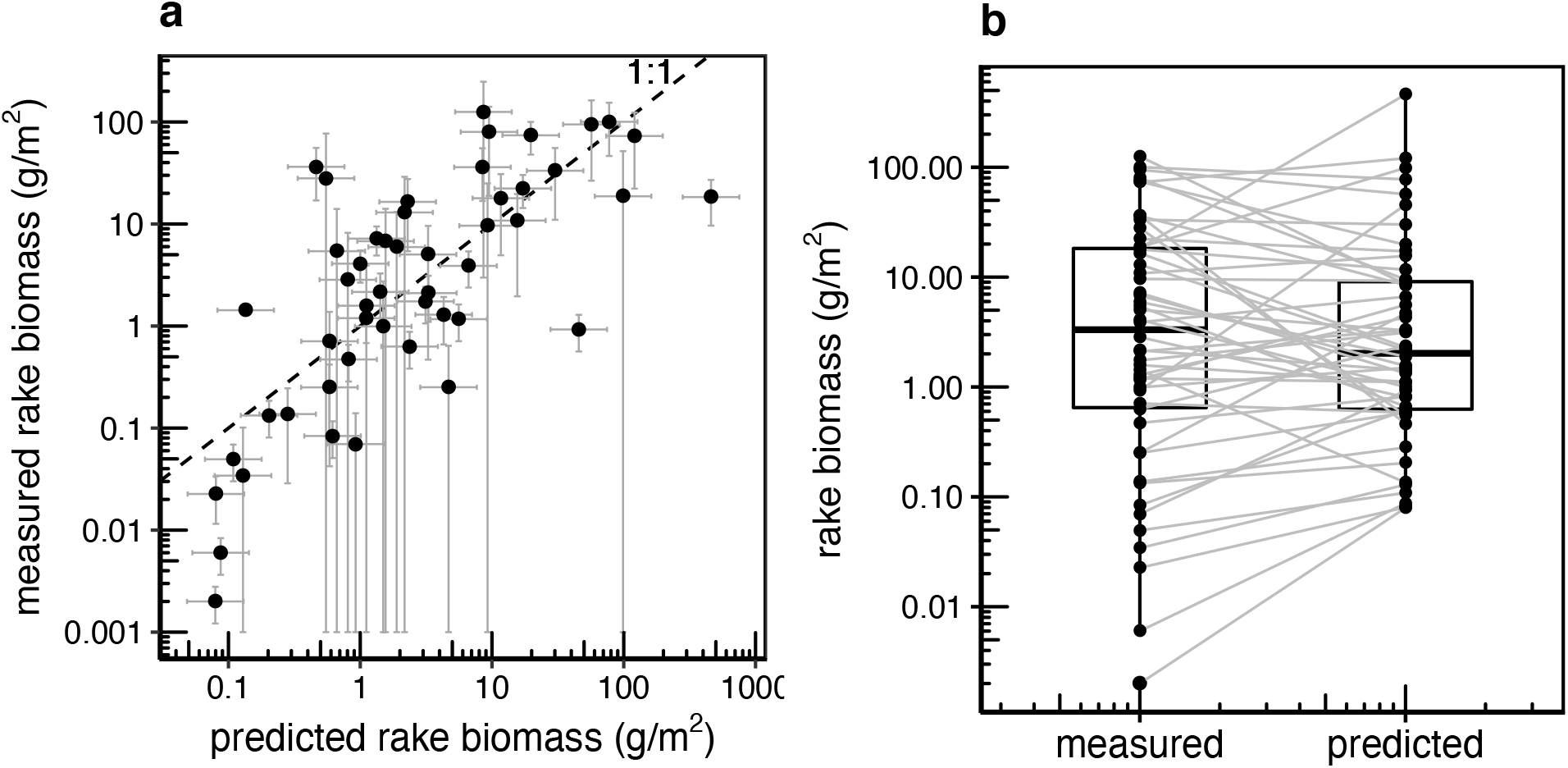
Evaluation of the PLSR cross-validation rake biomass predictions from echosounding. A) Scatterplot of measured vs predicted rake biomass, error bars represent the 95% confidence interval. B) Paired boxplot of measured vs predicted rake biomass, solid horizontal line within boxes represents the median, boxes extent the 25th and 75th percentiles and whiskers the 10th and 90th percentiles.

**Figure S7:**
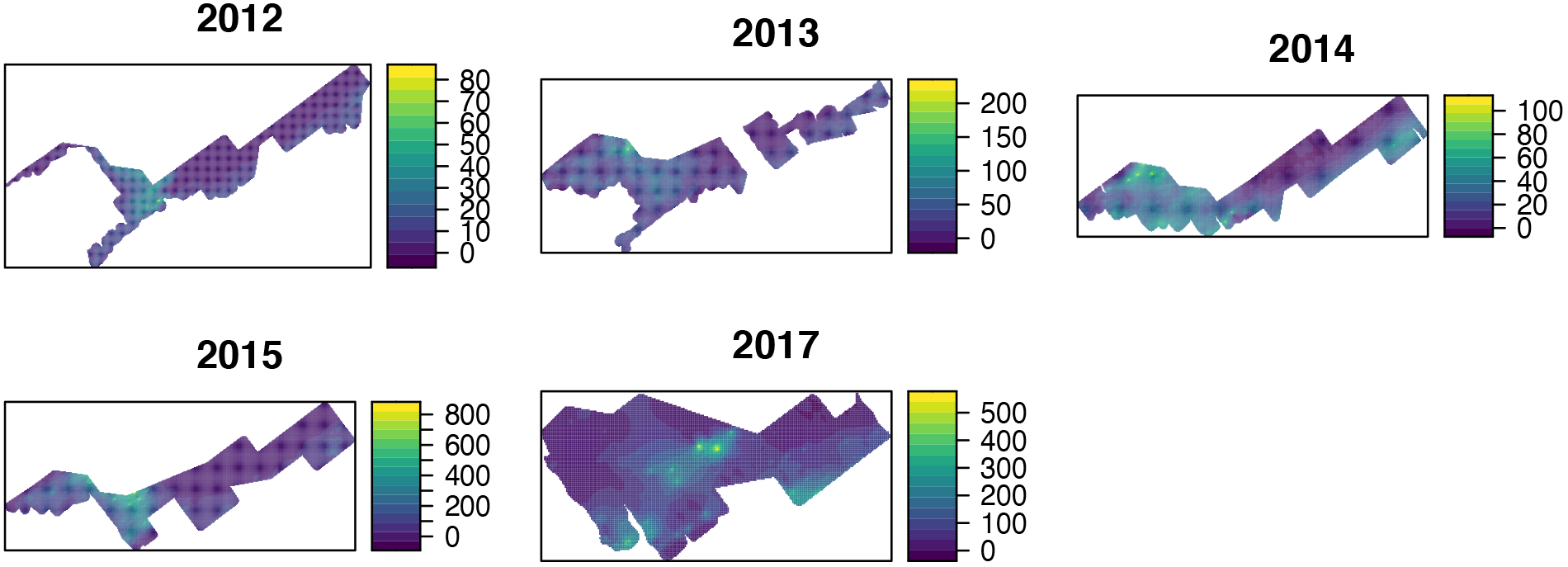
Kriging maps of biomass (g/m^2^) for whole-system estimation. Kriging surfaces are derived from two-step predictions of echosounding biovolume_10_.

**Figure S8:**
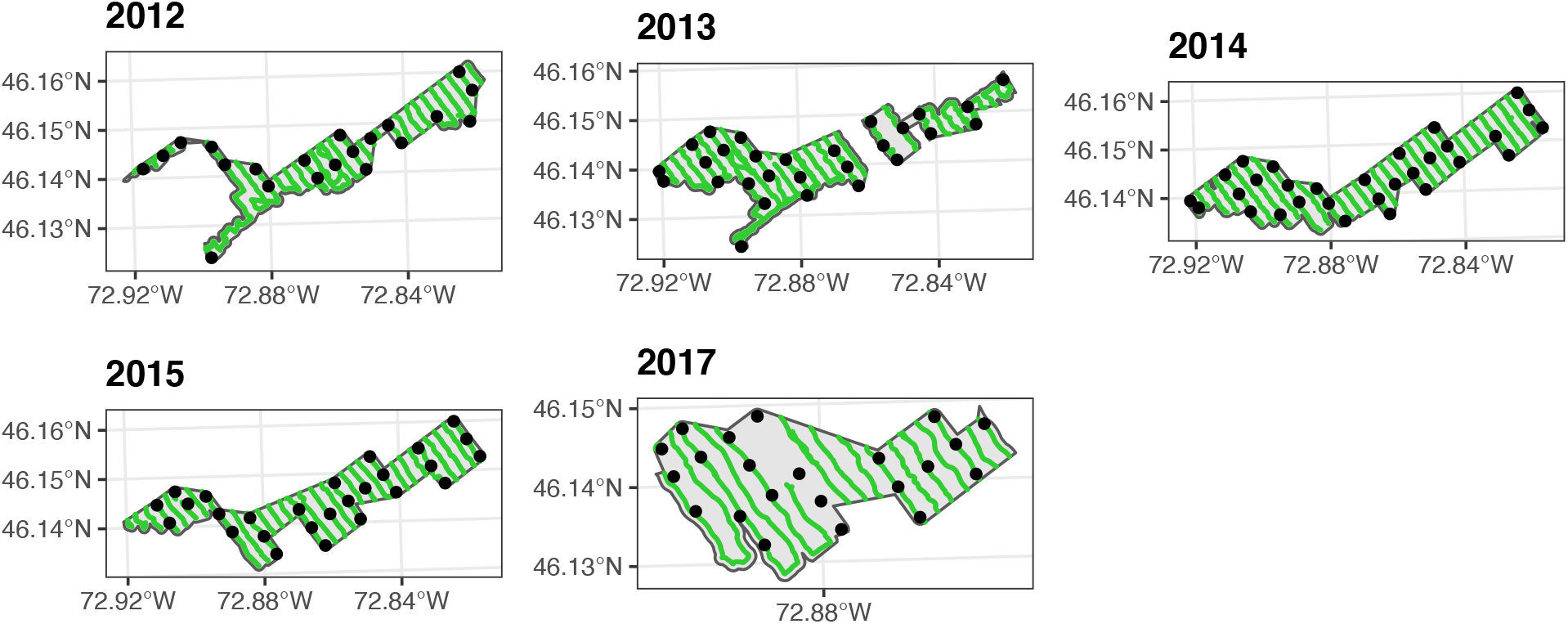
Location of rake samples (black dots) and of echosounding tracks (green lines) used for the whole-system SAV biomass estimations (2012-2017).

